# Human stem cell-derived GABAergic interneuron development reveals early emergence of subtype diversity followed by gradual electrochemical maturation

**DOI:** 10.1101/2024.12.03.626662

**Authors:** Marina Bershteyn, Hongjun Zhou, Luis Fuentealba, Chun Chen, Geetha Subramanyam, Daniel Cherkowsky, Juan Salvatierra, Meliz Sezan, Yves Maury, Steven Havlicek, Sonja Kriks, Seonok Lee, Michael Watson, Wai Au, Yuechen Qiu, Anastasia Nesterova, Derek Anderson, Brianna G Feld, Olga Kuzmenko, Maria Elena Grimmett, Victoria Hosford, Ji-Hye Jung, Tia Kowal, Alessandro Bulfone, Gautam Banik, Catherine Priest, Jorge Palop, Cory R. Nicholas

## Abstract

Medial ganglionic eminence-derived inhibitory GABAergic pallial interneurons (MGE-pINs) are essential regulators of cortical circuits; their dysfunction is associated with numerous neurological disorders. We developed human (h) MGE-pINs from pluripotent stem cells for the treatment of drug-resistant epilepsy. Here, we analyzed xenografted hMGE-pINs over the lifespan of host mice using single nuclei RNA sequencing. Comparative transcriptomics against endogenous human brain datasets revealed that 97% of grafted cells developed into somatostatin (SST) and parvalbumin (PVALB) subtypes, including populations that exhibit selective vulnerability in Alzheimer’s disease. Transplanted hMGE-pINs demonstrated rapid emergence of subclass features, progressing through distinct transcriptional states sequentially involving neuronal migration, synapse organization, and membrane maturation. We present molecular, electrophysiological, and morphological data that collectively confirm the derivation of diverse *bona-fide* human SST and PVALB subtypes, providing a high-fidelity model to study human MGE-pIN development and functional maturation as well as a compositional atlas for regenerative cell therapy applications.

## INTRODUCTION

GABAergic pallial interneurons (pINs) are a critical class of neuronal cells within the mammalian neocortex and hippocampus, where they play a key role in the development and regulation of cortical circuits. Changes in the proportion or functionality of interneurons have been linked to severe neurological disorders. Recent single cell and single nuclei RNA sequencing (sc/snRNAseq), patch sequencing, and spatial transcriptomics^1–6^ studies have catalogued the diverse molecular and functional properties of pINs in the endogenous mouse and human brain. Subsequent work has led to a unified cross-species cellular taxonomy, with pINs representing a broad class subdivided into major subclasses based on developmental origin^2,4,7–10^. The medial ganglionic eminence (MGE) generates more than half of the total pIN population, including PVALB, SST, SST/CHODL and LAMP5/LHX6 subclasses, while the caudal ganglionic eminence (CGE) generates other pINs, including VIP, SNCG, LAMP5 and PAX6 subclasses, with additional minor contributions from the preoptic area (POA)^11–24^. These broad subclasses are further divided into more specific interneuron subtypes exhibiting distinct features, such as molecular profile, morphology, distribution, connectivity, and electrophysiological properties. Diversification of pINs with shared developmental origin is initiated at the progenitor stage but becomes apparent after the postmitotic interneurons migrate into the pallium and engage in local interactions^16,25–29^. However, our understanding of how much of the differentiation is guided by intrinsic cell programming versus environmental factors remains incomplete^30–34^. While molecular profiling across developmental stages in different species enables the reconstruction of lineage trajectories and provides a blueprint of cell stage- and cell type-dependent signatures^2,26,27,29,35–38^, it is difficult to fully disentangle environmental influences from intrinsic programs in intact biological systems. Moreover, the protracted developmental timeline of human cortical neurons^39–43^, and the limited accessibility of primary human brain tissue further complicate the study of the regulatory programs that determine human pIN subclass and subtype identity.

Advances in directed differentiation methods using human embryonic stem cells (hESCs) and induced pluripotent stem cell (hiPSCs) (collectively referred to as hPSCs) and xenotransplantation provide an experimentally tractable framework to address some of these questions and limitations. Such an integrated approach is critical for advancing regenerative cell therapies for central nervous system (CNS) disorders. MGE-derived SST and PVALB pIN subclasses are of particular clinical relevance, as their dysfunction can result in numerous neurodevelopmental and neurodegenerative disorders of the nervous system^44–57^.

When transplanted into mouse neocortex or hippocampus, MGE-derived interneurons can migrate through brain parenchyma, persist for the lifespan of the host, and functionally integrate with host circuits, increasing inhibitory tone^58–62^. This suggested a potential therapeutic strategy for the treatment of drug-resistant focal epilepsy and other disorders characterized by hyper-excitable neural circuits^63,64^, and inspired efforts to generate MGE interneurons from hPSCs^40,65–74^. Accordingly, we developed a method for deriving highly pure, postmitotic, migratory GABAergic MGE-pINs from hPSCs (hMGE-pINs) and detailed their pre-transplant molecular identity and therapeutic effects in a mouse model of chronic focal epilepsy^72^. Despite their potential, systematic assessment of hPSC-derived grafts over time at single-cell resolution has been lacking. Considering that human cell grafts represent dynamic, evolving biological entities, it is crucial to understand their cellular composition and long-term stability for developing safe, efficacious, and mechanistically defined therapeutic interventions.

In this study, we present extensive single-nuclei RNA sequencing (snRNAseq) analysis of xenotransplanted hMGE-pINs into the mouse cortex and leverage published human brain transcriptomic data to assess grafted cell identity, maturation, and composition throughout the lifespan of the host. Unbiased computational analyses reveal early subclass diversification of human MGE-pINs potentiated by an *in vivo* neocortical environment. Together with histological and electrophysiological characterizations, our data demonstrate that transplanted hPSC-derived cells recapitulate key molecular, cellular and functional features of endogenous MGE interneurons, including SST and fast-spiking PVALB pIN subtypes. This work establishes an experimentally tractable platform for systematic evaluation of human interneuron development and provides insights into the mechanisms guiding hMGE-pIN specification, maturation, and function.

## RESULTS

### hESC-derived MGE-pINs are synchronized at a postmitotic migratory stage prior to transplantation

Fourteen independent lots of hMGE-pINs were generated from hESCs, as described previously, harvested, sorted for ERBB4-expressing cells to increase pIN purity, and cryopreserved (Table 1)^72^. Expression of on-target GABAergic MGE pIN markers and potential off-target markers was examined post-thaw. On average, cell lots were ∼97% positive for on-target markers *GAD1*, LHX6 and ERBB4, and ∼90% positive for CXCR7, MAF, and MAFB (Supp. Fig. 1A). The MGE progenitor marker NKX2-1 was downregulated by this stage compared to day 14^72^, while the CGE/dorso-caudal MGE marker COUPTF2 was expressed weakly (Supp. Fig. 1B). LHX8, which is maintained in some MGE-derived subpallial neurons^22,75^, was detected in less than 5% of the cells (Supp. Fig. 1A, C). Importantly, we did not detect other progenitor (OTX2/OLIG2), proliferative (MKI67), or neuronal off-target markers (cholinergic—ISL1/CHAT; hypothalamic—NKX2.2; CGE—SP8) (Supp. Fig. 1A, B).

We profiled all fourteen lots using scRNAseq to assess their reproducibility, purity, and cellular composition. Broad and uniform expression of neuronal (*DCX, MAPT, MAP2*), GABAergic (*GAD1, GAD2*), GE (*ARX, DLX1/2/5*) and MGE (*LHX6, SOX6*) markers (Supp. Fig. 1D, E, H) indicated a homogeneous population, with minor transcriptional differences between clusters 0-8. Seven out of nine clusters were enriched for more specific pallial markers (*ZEB2, MAF, MAFB, NXPH1, CXCR4, ERBB4*)^76–79^ and were designated accordingly “LHX6/MAF/ZEB2” clusters (Supp. Fig. 1E-H). This population likely represents the initial class of newborn postmitotic MGE-pINs, consistent with the MAF-expressing neurons present in the mouse E13.5 MGE subventricular zone (SVZ)^76,77^ (Supp. Fig. 1I). Cluster 5 was enriched for MGE-derived GABAergic inhibitory neuron markers (*SST, PDE1A* and *NPY*)^21,22,80,81^, and was designated “LHX6/SST/NPY” (Supp. Fig. 1D-J).

Finally, cluster 8 was enriched for MGE-derived pallidal mantle zone (PMZ) GABAergic inhibitory neuron markers (*LHX8, ENC1, ZFHX4, TSHZ2*) that regulate differentiation of globus pallidus and/or striatal populations^22,26,82–84^, and was designated “LHX6/ENC1/LHX8” (Supp. Fig. 1D-J). On average, cell lots consisted of ∼93% LHX6/MAF/ZEB2, ∼6% LHX6/SST/NPY, and ∼1% LHX6/ENC1/LHX8 (Supp. Fig. 1K). Importantly, markers corresponding to non-MGE off-target cell types were not expressed in any of the clusters or cell lots (Supp. Fig. 1E, G).

Next, we evaluated how our hMGE-pINs compare to endogenous populations in the human developing ganglionic eminences (GEs), including MGE, CGE, and lateral GE (LGE) progenitors and neurons, and other neuronal and glial populations^35^. Bioinformatic comparison (or prediction model)^72,85,86^ of transcriptomic similarity between the query and reference cell types demonstrated that both the LHX6/MAF/ZEB2 and LHX6/SST/NPY populations were postmitotic interneurons of pallial MGE identity and together comprised ∼98% of cells across lots (Supp. Fig. 2B, C). Similar results were obtained when comparing to a developing macaque GE reference dataset^37^ (data not shown).

To assess how our hMGE-pINs compare with other published GABAergic neurons generated from hPSCs, we analyzed available scRNAseq data representing diverse approaches, including directed differentiation using dual SMAD and WNT inhibition followed by SHH activation (“3i” method)^65,87,88^, transcription factor-mediated reprogramming using ASCL1 and DLX2 (“TF (AD)” method)^88,89^, and fused organoids with or without SHH activation (“CTX-GE” method)^90^ (Supp. Fig. 2A). Of these, the “3i” is most similar to our differentiation protocol^72^, yielding some MGE-pINs characterized by co-expression of *LHX6, ZEB2, MAF, MAFB* and *ERBB4* (Supp. Fig. 2D). However, this preparation also contained prominent populations of subpallial-like cells and proliferative progenitors (Supp. Fig.2 C, D). Cells generated by the “TF (AD)” method expressed neuronal markers (*DCX, STMN2*) and *SST*, but largely lacked MGE or pIN markers (*LHX6, MAF*) (Supp. Fig. 2D). Similarly, cells generated by fusing the dorsal and ventral organoids (“CTX-GE” method) generated a small population of *GAD2/SST*-expressing cells, which were otherwise devoid of MGE markers (*LHX6, MAF*), and instead co-expressed CGE (*SP8*) and cholinergic (*ISL1*) markers, suggesting a mixed identity (Supp. Fig. 2D). When compared to the endogenous human developing GEs, all the methods outlined above exhibited heterogeneous cell populations (Supp. Fig. 2B, C).

Next, we evaluated the developmental stage of the *in vitro*-derived cells by comparing their single-cell transcriptomes to those of endogenous cells across gestational stages GW9 – GW18^35^. Approximately (∼90%) of cells from our lots matched mid-second trimester (GW18) – a developmental stage that coincides with the transition from neurogenesis to neuronal migration during human cortical development^91^. This is consistent with the highly migratory behavior of postmitotic hMGE-pINs observed *in vitro* and *in vivo*^72^ (Supp. Fig. 3). Other published methods appeared to have less synchronized and less mature populations (Supp. Fig. 2E-G).

Finally, we measured spontaneous neuronal activity of hMGE-pINs, characteristic of immature migratory pINs during normal development^91^. Previously we demonstrated that hMGE-pINs release GABA prior to transplantation, which can be further potentiated by chemical depolarization^72^. Here, we transduced hMGE- pINs with calcium indicators and cultured them *in vitro* on primary mouse astrocytes for a few weeks. Spontaneous calcium transients were detected by 17 days in vitro (DIV), and were abolished by 1μM tetrodotoxin (TTX), a voltage-gated sodium channel blocker, but not by GABAR or AMPA/NMDAR blockers (Supp. Fig. 1L). These results indicate that early spontaneous neuronal activity in immature hMGE-pINs is mediated by intrinsic voltage-gated sodium-driven depolarizations rather than chemical synaptic inputs.

### Stable graft composition of xenotransplanted hMGE-pINs comprising PVALB and SST subclasses

To evaluate the migration, persistence and identity of hMGE-pINs *in vivo*, we transplanted neonatal immunocompromised mice (P1-P2) with a single deposit per cortical hemisphere and analyzed engrafted human cells using immunohistochemistry (IHC) (Supp. Fig. 3A, B). After the initial decline during the first few months, ∼5-10% of grafted human cells persisted for the lifespan of the host, maintaining high LHX6 expression (Supp. Fig. 3C). Transplanted cells migrated extensively by 1 MPT in both medial-lateral and rostro-caudal directions (>2.4mm), leaving no dense cores or cell clusters at or near the original deposit sites (Supp. Fig. 3B, C). Cell distribution remained unchanged from 1 to 4 MPT (Supp. Fig. 3C), indicating that interneuron migration was completed by 1 MPT. Consistently, interneuron morphology changed from a bipolar shape, typical of migratory doublecortin (DCX)-positive cells at 1 MPT to a multipolar morphology accompanied by increased expression of human-specific presynaptic marker synaptophysin (hSYP) (Supp. Fig. 3D-F).

To understand the fate of hMGE-pINs in the mouse brain, multiple lots were transplanted into the neonate cortex followed by dissections at 1, 3-4, 6-7, 12, and 18 MPT for snRNAseq (Fig. 1A). Total nuclei from dissected mouse cortices were extracted and stained with an antibody against human-specific nuclear antigen to enrich human cells using fluorescence-activated nuclear sorting (FANS) (Fig. 1A). Sequencing data were further bioinformatically sorted to remove any mouse reads from downstream analyses. After quality controls, the dataset comprised 130,557 transplanted human cells across five time points (2-4 independent lots per time point), with a median of 9,461 cells per sample (Table 1, Fig. 1B).

**Figure 1.**
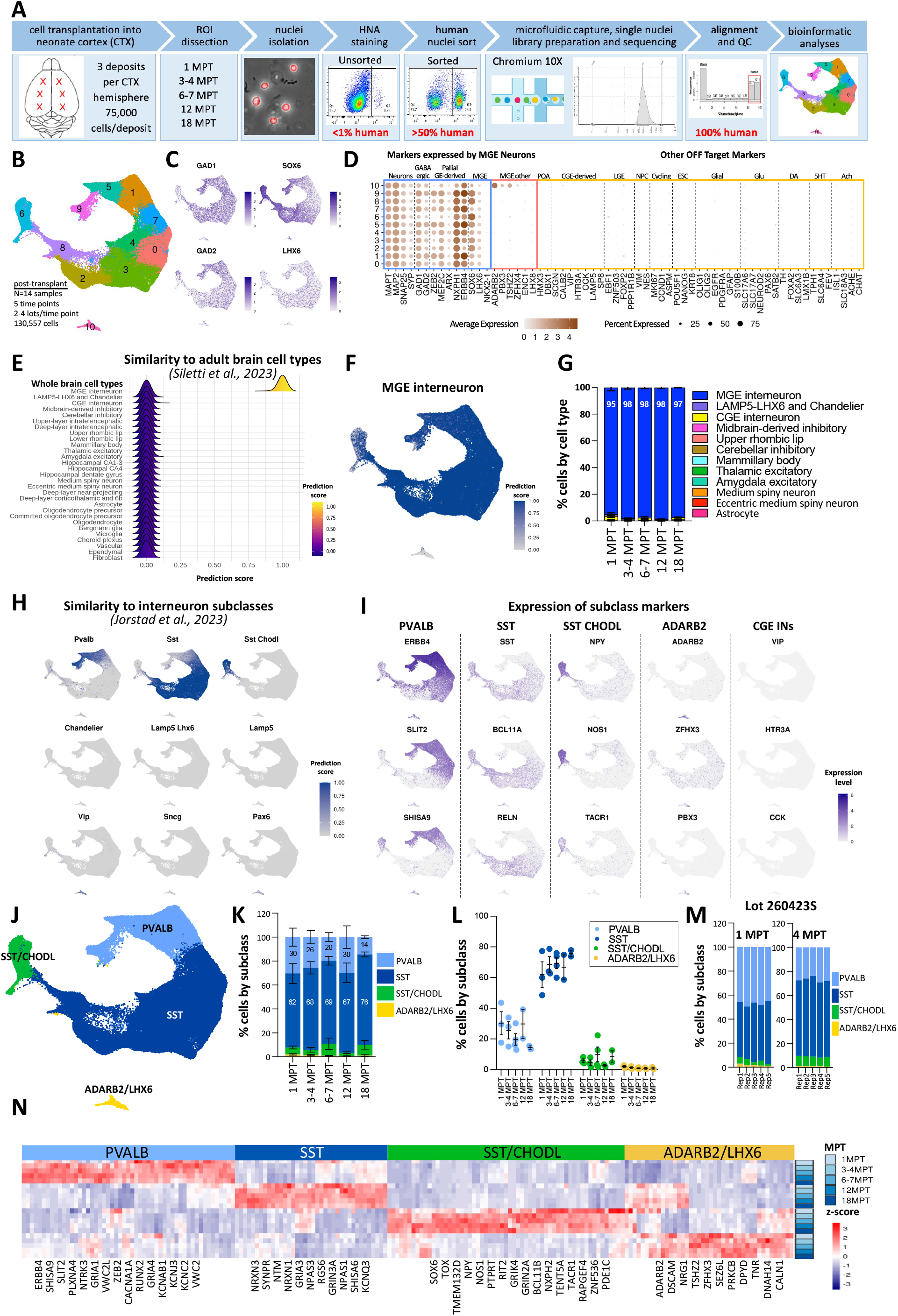
Graft composition from 1 to 18 MPT is stably comprised of PVALB and SST IN subclasses. (A) Experimental design. Cell lots (N=5) were transplanted into the neonate cortex in 3 deposits per hemisphere (75,000 cells/deposit). Cortical tissues (region of interest, ROI) were dissected at 1 MPT (N=3 lots), 3-4 MPT (N=3 lots), 6-7 MPT (N=4 lots), 12 MPT (N=2 lots) and 18 MPT (N=2 lots). Tissues from 2-3 animals were dissected for each lot and time point and pooled for nuclei isolation. Nuclei were stained with an antibody against human nuclear antigen (HNA), followed by fluorescence activated nuclear sorting (FANS) to enrich rare human nuclei. Sequencing reads were aligned to both human and mouse transcriptomes, filtering out any captured mouse reads from downstream analyses. (B) Integrated UMAP clustering of all human cells across time points. (C) Expression of GABAergic and MGE markers. (D) Expression of genes that characterize diverse cell categories by clusters, including general markers of neurons, GABAergic and MGE neurons, preoptic area (POA), caudal and lateral GE (CGE/LGE), neuronal progenitors (NPC), cycling cells, ESCs, as well as genes associated with glial cells, glutamatergic neurons (Glu), dopaminergic neurons (DA), serotonergic neurons (5HT) and cholinergic neurons (Ach). (E) Cell type prediction scores comparing transplanted cells with human adult brain superclusters of diverse brain cell types^92^. (F) Prediction scores of MGE interneuron fate for each transplanted cell. (G) Graft composition over time based on predicted adult brain superclusters. (H) Prediction scores of MGE- and CGE-derived interneurons corresponding to human interneuron subclasses in the middle temporal gyrus (MTG)^10^. (I) Expression of ADARB2-enriched genes consistent with MGE GABAergic subpallial neurons^22^. (J) Expression of markers associated with pIN subclasses listed at the top of each column. (K) Assignment of interneuron subclass identity. (L, M) Graft composition over time based on interneuron subclasses. (N) Reproducibility of cell composition at 1 and 4 MPT for the same lot transplanted from different vials (n=5 technical replicates). (O) Expression of top subclass- enriched genes. (G, K, L) Data are shown as mean ± SEM.

Using Seurat integration, we identified 11 high-quality clusters, without notable batch effects between samples (Fig. 1B; Supp. Fig. 4A-C). These clusters consistently expressed neuronal, GABAergic, pallial MGE-derived markers, while off-target markers were not observed over the 18 MPT period (Fig. 1C, D; Supp. Fig. 4D). To interpret grafted cell identity, we compared the nuclear transcriptomes of the hESC-derived cells to a diverse reference snRNAseq dataset from adult human brain, comprising 30 major cell classes^92^. Prediction analysis^85,86^ indicated that overall, more than 97% of transplanted cells corresponded specifically to canonical MGE interneurons (Fig. 1E-G; Supp. Fig. 4E). Moreover, similarity scores compared to the various human brain regions indicated that grafted cells had higher correspondence to telencephalic regions, starting with cerebral cortex, followed by hippocampus and amygdala, with low prediction scores (<0.25) corresponding to the basal forebrain. There was no similarity to diencephalon, midbrain, hindbrain, or spinal cord (Supp. Fig. 4F).

For higher resolution of interneuron subclass identity, we compared xenografted hMGE-pINs against multiple human adult reference datasets with the most recent annotations of interneuron subclasses and subtypes from different cortical regions, including the middle temporal gyrus (MTG), primary motor cortex (M1), and prefrontal cortex (PFC)^10,36,93,94^. Regardless of the reference dataset employed, prediction analyses yielded consistent results, demonstrating that transplanted hMGE-pINs comprised PVALB, SST and SST/CHODL subclasses (Fig. 1H illustrates comparison to MTG; Fig. 2H illustrates comparison to PFC; Supp. Fig. 4G illustrates comparisons to MTG and M1). Remarkably, subclass prediction was robust already by 1 MPT, long before the cells reached full functional and neurochemical maturation, with slight increases in scores over time (Supp. Fig. 4G). Notably, there was no transcriptional similarity to the MGE-derived chandelier or LAMP5/LHX6 neurogliaform cells, nor to the CGE- or LGE-derived subclasses (Fig. 1E, H).

**Figure 2.**
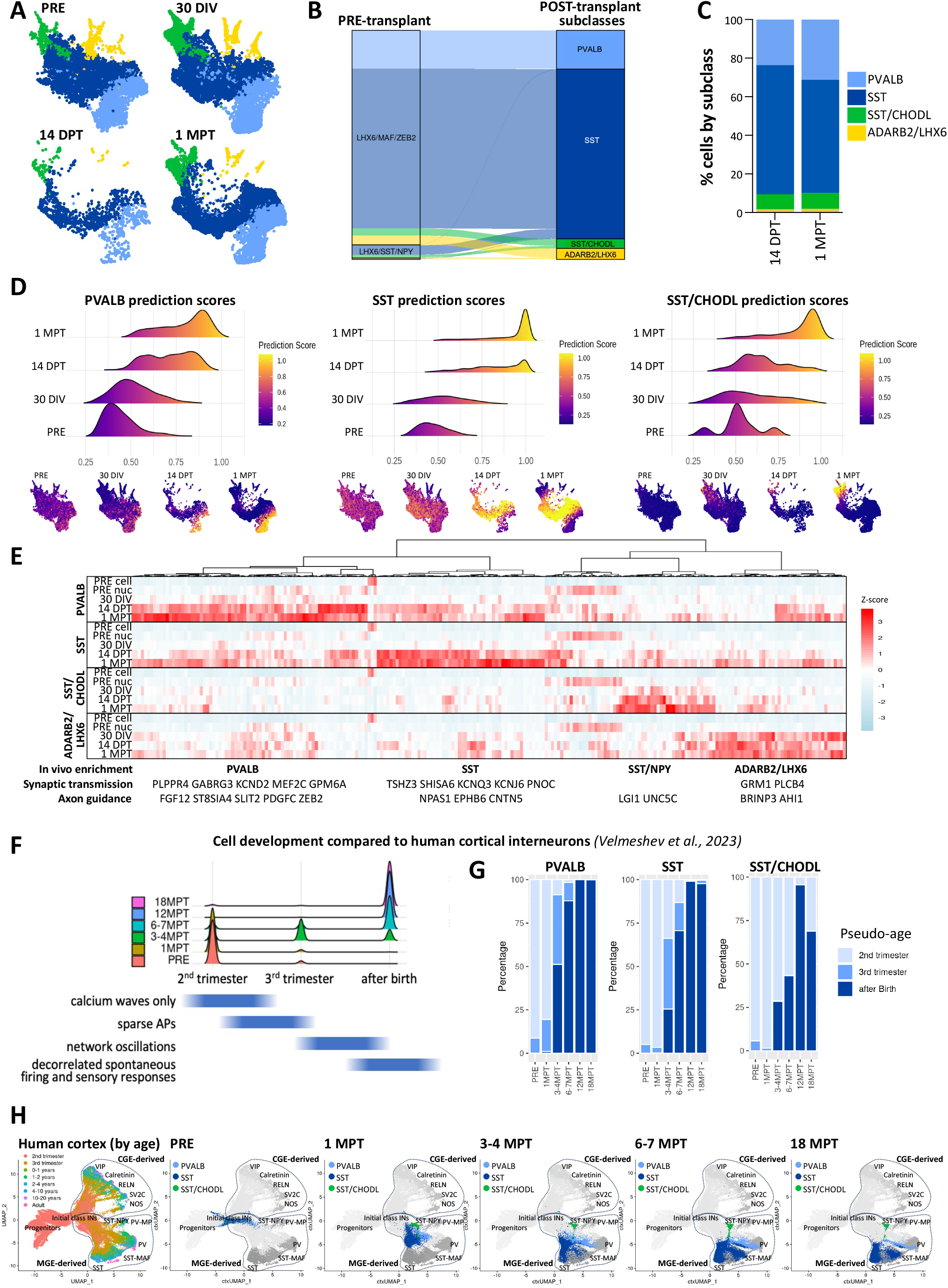
Xenotransplantation environment triggers expression of synaptic genes, promoting cell fate differentiation. (A) Integrated cell clustering of one lot (010519S1) sequenced prior to transplantation (PRE), after one month of *in vitro* culture (30 DIV), 14 days post-transplantation (14 DPT) and 1 MPT, colored by subclass. (B) Transcriptomic relationship between pre- and post-transplant clusters. (C) Graft composition by subclass at 14 DPT and 1 MPT. (D) Prediction scores for PVALB, SST and SST/CHODL subclasses across samples compared to the adult MTG reference shown as a density distribution (top) and feature plots (bottom). (E) Expression of subclass-specific genes upregulated in 1 MPT vs 30 DIV samples. (F-H) Evaluation of graft development post-transplantation compared with human cortical development spanning from prenatal to postnatal stages. The quantification is based on predicted cell stage analysis using cortical developmental datasets^36^ as a reference. (F) Distribution of grafted cell maturation states over time relative to broad human stages indicated below the ridge plots, corresponding to developmental changes in neuronal physiology (adapted from *Wallace, Pollen 2024*)^43^. (G) Grafted cell maturation states over time for each of the three main subclasses. (H) The re-clustering of endogenous prenatal and postnatal human CGE- and MGE-derived interneurons from prefrontal cortex (PFC)^36^ (left), and hESC-derived pIN at different time points projected onto human PFC interneuron development trajectories (right).

Interestingly, a single cluster (#10), representing less than 3% of the transplanted cells, did not correspond to canonical MGE interneurons (Fig. 1F). This cluster co-expressed *GAD1, GAD2*, *SOX6,* and *LHX6* (Fig. 1C, D), together with *ADARB2*, *LHX8, GBX1,* and other markers (Fig. 1D, I, J) consistent with the MGE-derived subpallial Lhx8 Gaba subclass identified in the cerebral nuclei by van Velthoven et al., 2024 (subclass #057: NDB-SI-MA-STRv)^22^. Markers of other potential MGE lineages, including striatal GABAergic interneurons and pallidal dual GABAergic-cholinergic neurons (*ST18, PROX1, CHAT, ACHE*)^22^ or the MGE-derived neurogliaform cells (*LAMP5*)^6,77–80^ were not expressed (Fig. 1D and data not shown), suggesting that cluster 10, henceforth designated “ADARB2/LHX6” (Fig. 1K), likely represents subpallial MGE-derived GABAergic neurons that are not normally found in the cortex. No CGE-specific markers (*SP8, SCGN, VIP, HTR3A, CCK*) were detected in any of the transplanted human clusters (Fig. 1D, J).

Gene expression patterns of known subclass markers corroborated the results from the prediction model. PVALB lineage markers, including *ERBB4, SLIT2, SHISA9, RUNX2* (Fig. 1J, O; Supp. Fig. 4H) alongside potassium channel subunits genes linked to fast-spiking properties (*KCNJ3, KCNAB1, KCNC2*)^21^ (Fig. 1O; Table 2) were enriched in the predicted PVALB population, although *PVALB* itself was not yet expressed. Likewise, SST lineage markers, including *SST, RELN, BCL11A, NPAS1, NPAS3, SYNPR, GRIN3A,* and *GRIA3* were enriched in the predicted SST population (Fig. 1J, O; Table 2), while the predicted SST/CHODL population expressed *SST* along with *NPY, NOS1*, *TACR1, CHODL, and TOX* (Fig. 1J, O; Table 2).

Based on highly concordant results from the data-driven clustering analysis, marker expression patterns and prediction model, we determined that hMGE-pIN grafts consisted of 20-40% PVALB, 60-80% SST, 5-10% SST/CHODL and <3% ADARB2/LHX6 subclasses (Fig. 1K-M). The proportions remained generally stable over time (Fig. 1L, M), although there was an initial decrease in PVALB cells from 1 to 4 MPT (Fig. 1N). Independent assessments at 1 MPT and 4 MPT using five separate frozen vials from the same lot showed that variation between technical replicates was minimal (Fig. 1N), indicating that a single measurement provides reliable representation of graft composition for a given lot. Importantly, there was no emergence of new or off-target cell types from the grafts during the 18 months after transplantation.

To validate the profiling results in tissue sections, we performed fluorescence in situ hybridization (FISH) with probes designed to distinguish the main cell populations. Using human-specific *MALAT1* probe, we confirmed that nearly all the grafted cells expressed the pan-GABAergic marker *GAD1* (Supp. Fig. 4I-L). More specific subclass markers *SLIT2* and *SST* are expressed in complementary, partially overlapping gradients in endogenous (Supp. Fig. 4H) and hESC-derived PVALB and SST pINs, respectively (Fig. 1J). Therefore, to identify the putative PVALB population, we quantified *SLIT2+/SST-* cells (Supp. Fig. 4K, L), whereas to identify the putative SST population, we quantified all *SST+* cells. Since *SST* expression increases over time in both the SST and SST/CHODL subclasses (Supp. Fig. 4J), the fraction of *SST*-type cells was underestimated at 1 MPT, increasing ∼6-fold by 4-18 MPT (Supp. Fig. 4L). In contrast, the proportions of *SLIT2*+ and *NPY*+ cells yielded similar estimates to the fractions of PVALB and SST/CHODL subclasses based on snRNAseq, at early and late time points (Supp. Fig. 4KL; Fig. 1M). Overall, while the FISH-based experiments provided an *in-situ* validation of snRNAseq results, they underscored the limitations of the probe-based approach relying on just a few markers for quantitative assessment of cell types, especially at early post-transplant stages.

### Transplantation of hMGE-pINs into the mouse neocortex promotes subclass differentiation

One of the fundamental questions in interneuron biology is understanding the earliest determinants of cell fate. We explored whether subclass identity of hMGE-pINs could be discerned *in vitro* after extended culture, using the computational prediction model, and if subclass identity could be predicted *in vivo* even sooner than 1 MPT—thereby separating the effects of timing and environment. To investigate this, we sequenced nuclei from one lot (010519S1) either immediately post-thaw, after 30 days of *in vitro* co-culture with astrocytes (30 DIV), or 14 days post-transplantation (14 DPT), integrating these data with 1 MPT snRNAseq data from the same lot (Supp. Fig. 5A, B). Clustering analysis enabled assignment of subclass identity to the pre-transplant cells based on transcriptomic similarities between pre- and post-transplant populations (Fig. 2A; Supp. Fig. A, B). Alluvial plots analysis suggested that the main pre-transplant LHX6/MAF/ZEB2 population gives rise to all four subclasses observed *in vivo*, whereas the LHX6/ENC1/LHX8 population contributes only to the ADARB2/LHX6 subclass (Fig. 2B). Notably, PVALB cells are almost exclusively derived from the LHX6/MAF/ZEB2 pre-transplant population, whereas a small fraction of SST and SST/CHODL subclass cells appeared related to the pre-transplant LHX6/SST/NPY population (Fig. 2B).

While the transcriptomic relationship between the pre- and post-transplant populations could be established, similarity between the pre-transplant samples and the human adult endogenous interneurons was weak, as evidenced by the low prediction scores, even after 30 DIV (Fig. 2D). Moreover, markers such as *ERBB4* (PVALB-enriched) and *RELN* (SST-enriched) did not exhibit subclass-specific enrichment after 30 DIV that is observed *in vivo* (Supp. Fig. 5C; Fig. 1J). In contrast, 14 DPT was similar to 1 MPT in terms of clustering, composition and prediction results (Fig. 2A-D), with subclass-specific gene expression patterns already emerging by 14 DPT (Fig. 2E; Table 3). Consistently, differential gene expression (DGE) analysis between 30 DIV and 1 MPT samples revealed enrichment for biological processes involved in synaptic transmission and axon guidance in the transplanted cells (Supp. Fig. 5D). These results suggest that, while subclass identity is likely predetermined in newborn pINs, the post-transplant neocortical environment promotes their differentiation by triggering transcriptional programs that support cell type-specific interactions, revealing subclass fate when human interneurons are still immature.

### Interneuron development is characterized by major transcriptional changes during the first 3 MPT, followed by gradual unfolding of gene modules involved in regulation of synaptic transmission and membrane potential

To understand the maturation of ESC-derived hMGE-pINs in the context of human interneuron development, we utilized as a reference an integrated human dataset containing cortical interneurons spanning from the second trimester of gestation to adulthood^36,94–96^. Given that the impact of species-specific experience and activity-dependent effects on human interneuron maturation cannot be modeled accurately in a xenotransplantation environment ^31,33,97,98^, we narrowed the assessment to broad developmental stages corresponding to major physiological changes in circuit development^43^ (Fig. 2F). When projected onto human cortical interneuron developmental trajectories, pre-transplant cells (PRE) aligned with the initial class of endogenous postmitotic interneurons corresponding to the 2^nd^ trimester stage (Fig. 2G, H), consistent with previous analysis using only the developing human GE reference^35^ (Supp. Fig. 2E-G). At 1 MPT, cells shifted along the MGE developmental trajectory – and not CGE, with clear subclass separation, although most cells still matched the 2nd trimester (Fig. 2F-H). By 3-4 MPT, over half of the PVALB and SST cells corresponded to the 3rd trimester or later stages. This trend continued, with cells progressively advancing to early postnatal-like stages by 18 MPT (Fig. 2F-H), indicating continued *in vivo* maturation.

To explore the molecular programs underlying cell maturation, we pooled pre- and post-transplant samples, clustering them without integration. This analysis revealed three major transcriptional states: pre-transplant (PRE), 1 MPT, and 3+ MPT, as illustrated by shifts in the UMAP space (Fig. 3A-C). Across these major transitions, the early marker *DCX* was downregulated after transplantation, whereas neuronal (*MAP2*), GABAergic (*GAD1*, *GAD2*) and pan-MGE (*LHX6, SOX6*) markers remained stable (Fig. 3D). DGE analysis indicated that the PRE stage was enriched for genes involved in forebrain development, neuronal differentiation, cytoskeletal regulation/migration, and neuron projection/axon development (Fig. 3E, F; Supp. Fig. 4C-E). By 1 MPT, projection/axon guidance and neuronal migration programs were still active, but additional programs involved in cognition, learning and memory, synapse organization and glutamatergic transmission began to emerge (Fig. 3F; Supp. Fig. 4C-E). After 3 MPT, genes involved in cell adhesion, modulation of chemical synaptic transmission and regulation of ion transport and membrane potential became prominent (Fig. 3F; Supp. Fig. 4C-E). GO enrichment analysis for cellular components and molecular function confirmed the transitions from cytoskeletal programs pre-transplant to synaptic signaling activity at 1 MPT, and channel activity beyond 3 MPT (Supp. Fig. 4C-E), providing transcriptomic evidence for the kinetics of grafted hMGE-pIN maturation and circuit integration.

**Figure 3.**
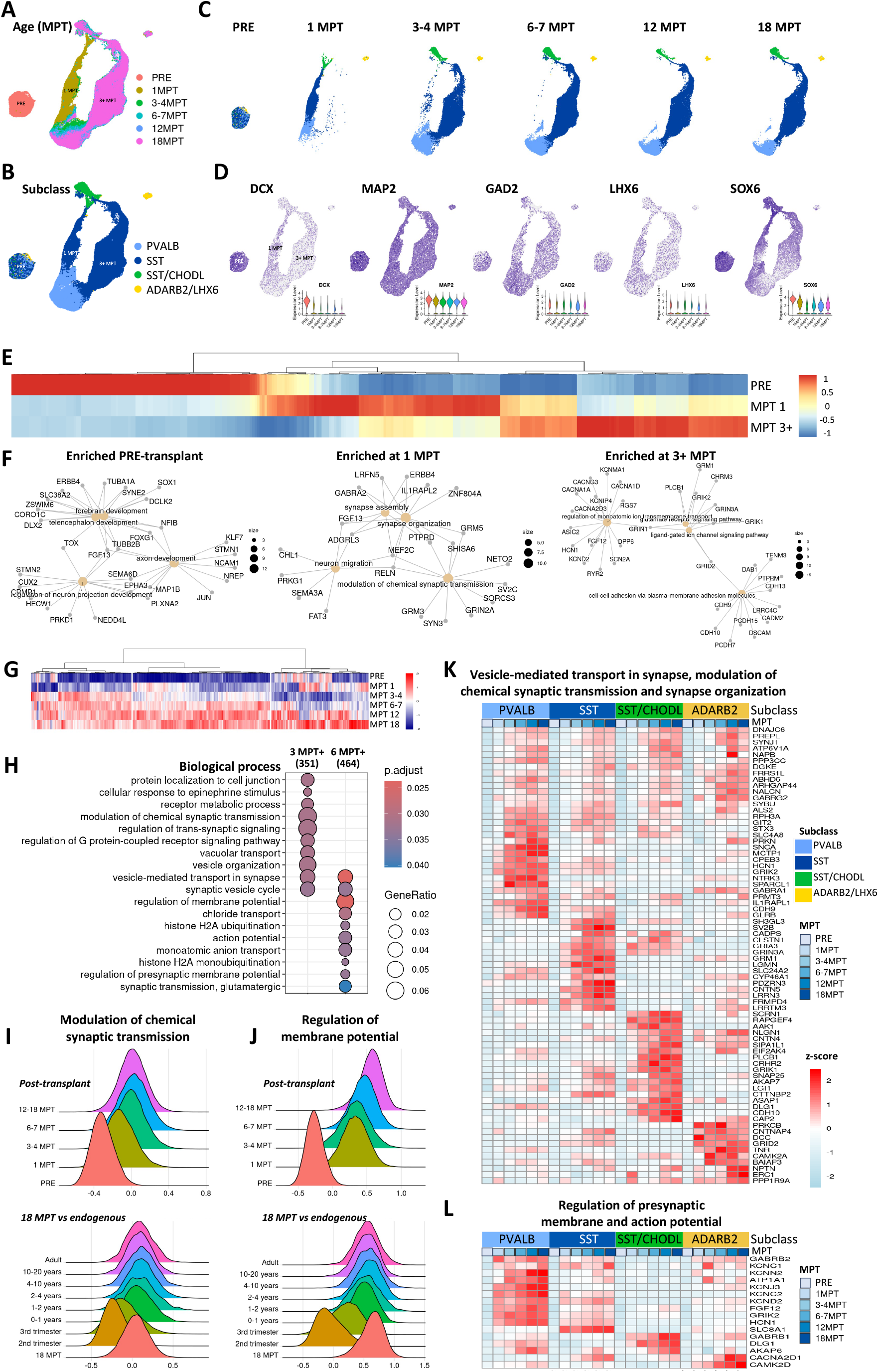
Cell development is characterized by major transcriptional changes during the first 3 MPT followed by gradual unfolding of gene modules involved in regulation of synaptic transmission and membrane potential. (A-C) Merged clustering analysis, without integration. Cells are colored by MPT (A), by subclass with all the stages combined (B), or separated by MPT (C). (D) Expression of neuronal, GABAergic and MGE markers across transcriptional states. (E) Differential gene expression analysis reveals three major transcriptional stages corresponding to pre-transplant (PRE), 1 MPT and 3-18 MPT. (F) Genes of top 4 GO biological processes enriched at each of the main transcriptional states. (G) Genes gradually upregulated over time. (H) Top GO biological processes enriched after 3 MPT and 6 MPT. (I, J) Expression of gene modules for chemical synaptic transmission (I) and membrane potential (J) in the grafts over time (top plots), as well as for 18 MPT grafts in the context of human cortical development (bottom plots). (K, L) Expression of cell type specific genes gradually upregulated over time and involved in vesicle-mediated transport in synapse, modulation of chemical synaptic transmission and synapse organization (K), as well as regulation of presynaptic membrane and action potential (L).

Beyond 3 MPT, the samples overlapped significantly in the UMAP space (Fig. 3A-C), suggesting that subsequent changes may primarily affect the levels rather than the types of genes that are expressed and/or posttranscriptional mechanisms that support interneuron maturation. Indeed, linear regression analysis to test the association of gene expression with cell age revealed gradual upregulation of genes involved in G protein-coupled receptor signaling, synaptic vesicle organization, ion transport, glutamatergic synaptic transmission and action potential regulation (Fig. 3G, H; Table 4). We examined temporal expression of the largest non-overlapping gene modules in grafted and endogenous cINs: “modulation of chemical synaptic transmission”, upregulated significantly after 3-4 MPT (Fig. 3H, I) and “regulation of membrane potential”, upregulated significantly after 6-7 MPT (Fig. 3H, J). Both modules showed continuous upregulation post-transplant (Fig. 3I, J top rows), with the oldest grafted sample (18 MPT) aligning with neonatal and postnatal ages (Fig. 3I, J bottom rows). A subset of genes contributing to these modules were broadly expressed, but many showed subclass-specific enrichment over time (Fig. 3K, L). For example, voltage-gated potassium channels encoding Kv3 (*KCNC1* and *KCNC2*), which are important for the sustained fast-spiking properties in PVALB INs^99^, exhibited enrichment with continuous post-transplant upregulation in the hESC-derived PVALB subclass (Fig. 3L), highlighting the progressive functional maturation of these cells *in vivo*.

### Age-dependent functional maturation of the action potential waveform and intrinsic membrane properties of transplanted hMGE-pINs

Previous analysis showed that hMGE-pINs remained electrophysiologically immature 7.5 MPT^72^. The delayed onset and gradual upregulation of the membrane potential-related genes observed in our transcriptomic studies prompted a more detailed examination at later stages. We conducted whole-cell patch-clamp recordings of transplanted GFP-expressing hMGE-pINs in acute coronal cortical slices from 12 to 20 MPT to assess their intrinsic membrane and action potential (AP) firing properties. Our recordings revealed progressive maturation in AP firing rates and AP kinetics (Figs. 4A, B), including shorter AP half-width, rise time, and afterhyperpolarization (AHP) delay (Figs. 4C–F), as well as larger AHP amplitudes (Fig. 4F). Together, these findings indicate that hMGE-pINs sustain long-term maturation of waveform over the entire lifespan of the host mouse (Fig. 4G). To test whether the transplanted hMGE-pINs integrated into local circuits and received local inputs, we recorded spontaneous postsynaptic currents (sPSCs) at the different timepoints. We observed an age-related increase in the frequency of sPSCs onto hMGE-pINs (Figs. 4H), indicating that intrinsic maturation and circuit integration of hMGE-pINs occur in parallel.

**Figure 4.**
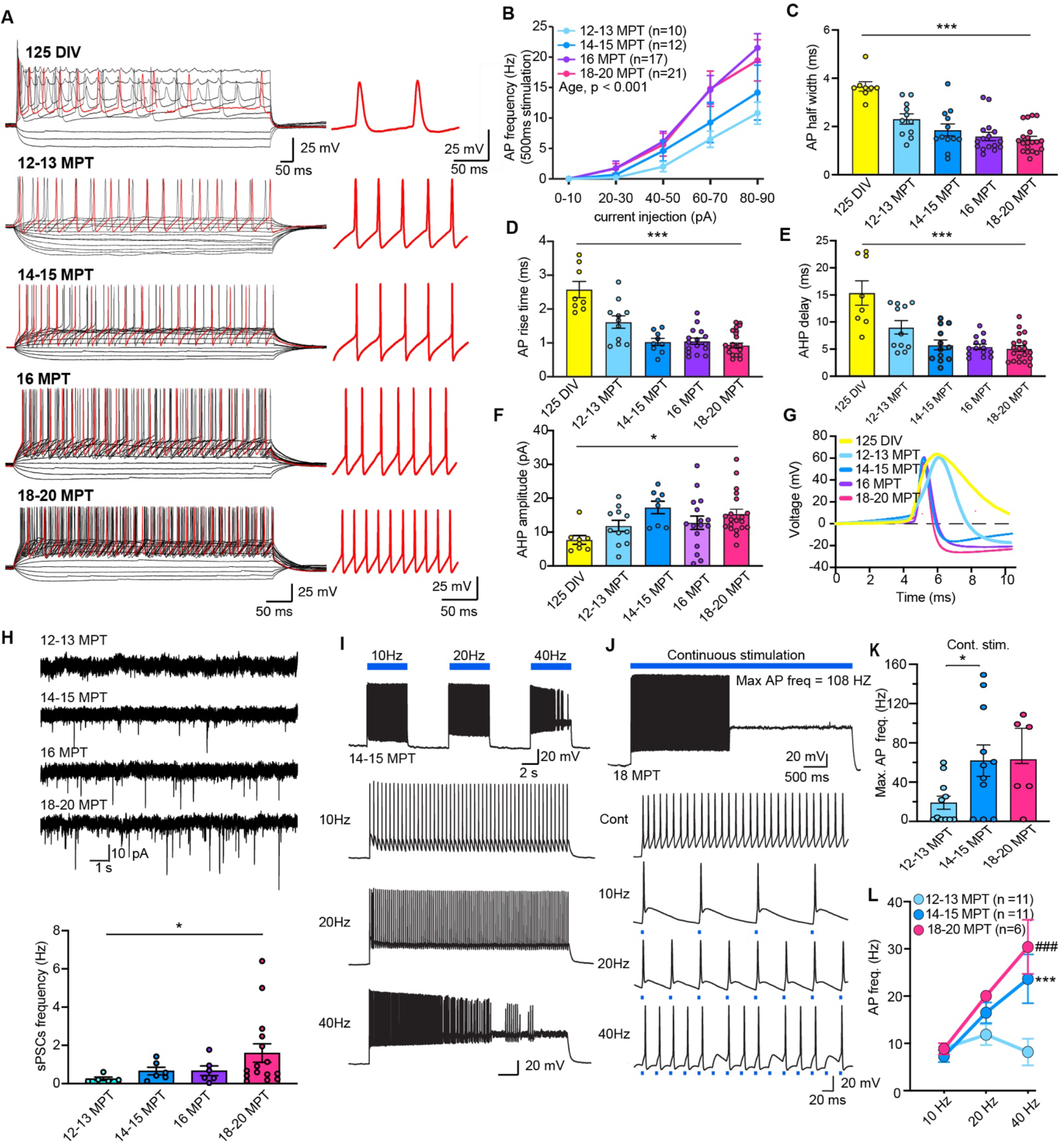
Action potential (AP) kinetics of developing hMGE-pINs after xenotransplantation into the mouse brain. Whole-cell patch-clamp recordings from GFP/ChR2-positive hMGE-pINs after 125 DIV culture (n=8) and in the cortex of acute coronal brain slices from 12-13 MPT (n=10), 14-15 MPT (n=12), 16 MPT (n=17) and 18-20 MPT (n=21). (A) Representative traces of AP firing from each age group. (B) Mean frequency of APs evoked by incremental current intensities for 500 ms. (C-F) Intrinsic properties of the AP waveform, including half width (C), rise time (D), fall time (E), and afterhyperpolarization amplitude (F) development over time. One-way ANOVA: ***, p<0.0001; *, p<0.05. (G) Traces of the averaged waveforms from each age group. (H) Spontaneous postsynaptic currents (sPSC) recorded from transplanted cells for each age group with representative raw traces (top) and average frequency (bottom). One way ANOVA: *, p<0.05. (I) Representative traces from CHR2-expressing hMGE-pINs at 14-15 MPT in response to 10, 20 and 40 Hz photo-stimulation. (J) Representative traces from the 18 MPT age group in response to continuous, 10 Hz, 20 Hz and 40 Hz photo- stimulation. The instantaneous maximum firing frequency in continuous trace is 108 Hz. (K) Maximum instantaneous frequency of the transplanted cells over time in response to 5 second continuous stimulation. One way ANOVA with post hoc analysis, * p<0.05; (L) Average AP frequency in response to the 1^st^ second of photo-stimulation at 10, 20 and 40 Hz. Two-way ANOVA with post hoc analysis, ***, 14-15 MPT vs. 12-13 MPT, p<0.0001; ###, 18-20 MPT vs. 12-13 MPT, p<0.0001. Data shown are mean ± SEM.

We also employed channelrhodopsin-mediated optogenetic stimulation to precisely elicit APs in transplanted hMGE-pINs, using 1 ms photo-stimulation at 10, 20, and 40 Hz (Figs. 4I, J) for 5 seconds. Human MGE-pINs consistently generated photo-stimulatory frequency-dependent APs at 10, 20, and 40 Hz, particularly after 14 MPT, indicating their ability to sustain high firing rates over extended periods of stimulation. Notably, we identified numerous hMGE-pINs capable of producing sustained APs at very high firing rates (>100 Hz for several seconds) under continuous optogenetic stimulation (Fig. 4K), demonstrating their capacity for fast-spiking activity. Additionally, and consistent with the above results, hMGE-pINs exhibited age-dependent increases in maximum firing rates, which rose from 20 Hz at 12–13 months to 60 Hz at 18–20 months, along with improved AP fidelity over time (Figs. 4K, L). These findings underscore the age-dependent maturation of hMGE-pINs and their ability to sustain high-frequency firing as they integrate into the host brain circuitry.

### Transplanted hMGE-pINs differentiated into non-adapting fast-spiking and adapting non-fast-spiking interneurons

We previously showed that murine transplanted MGE-derived interneurons developed into non-adapting fast-spiking (FS) and adapting non-fast-spiking (NFS) subtypes, acquiring the distinct electrophysiological properties of Pvalb and Sst interneurons, respectively^55^. To assess whether the older hESC-derived transplanted MGE-pINs (16–20 MPT) similarly differentiate into FS and NFS cells, we examined the action potential (AP) firing rates in response to increasing injected currents (0–150 pA). Our findings indicated that 64% of the recorded cells (23 out of 36 cells) exhibited little to no AP frequency saturation across current steps and high firing rates that increased proportionally with increasing current injection (Figs. 5A), consistent with the non-adapting fast-spiking phenotype of PVALB cells^4^. The remaining 36% of the cells (13 out of 36 cells) exhibited lower firing rates at higher current steps, indicative of depolarization block, which is a pattern typical of the adapting non-fast-spiking SST interneurons^4^. Notably, since we primarily recorded from deeper cortical layers (4–5) and larger cells, this likely explains the higher proportion of FS phenotypes, which are more commonly associated with PVALB cells in these layers^4^.

**Figure 5.**
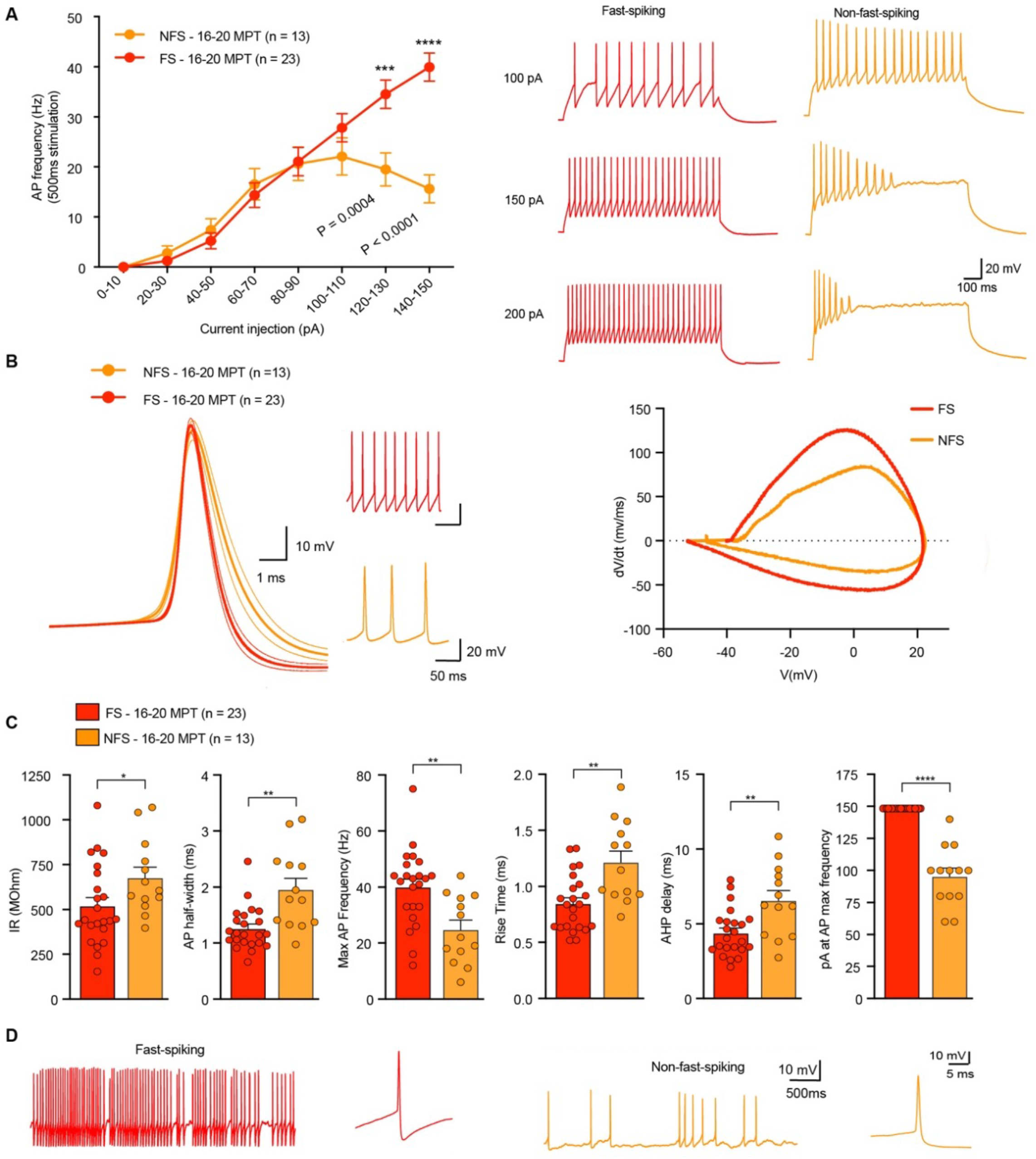
Transplanted hMGE-pINs differentiate into non-adapting fast-spiking (FS) and adapting non-fast-spiking (NFS) interneurons. (A) Left panel: mean frequency of AP evoked by incremental current intensities for 500 ms in transplanted hMGE-pINs demonstrating FS (n=23 cells, from 5 mice) and NFS (n=13 cells, from 5 mice) phenotypes. Right panels: representative traces of AP firing in response to 100, 150 and 200 pA current injection recorded from FS and NFS hMGE-pIN groups. (B) Left panel: averaged waveforms from FS and NFS hMGE-pIN cell groups with example APs from shown. Dotted lines indicate standard error of the mean. Right panel: averaged phase plot of FS and NFS hMGE-pINs. (C) Intrinsic properties of the AP waveform from FS and NFS hMGE-pIN cell groups at 16-20 MPT, including membrane resistance, AP half width, maximum instantaneous AP frequency, AP rise time, and AHP delay. Data shown are mean ± SEM. Unpaired two-tailed student t-test: * p<0.05, ** p<0.01, **** p<0.0001. (D) Representative traces of spontaneous AP firing at resting membrane potential from FS and NFS hMGE-pIN cell groups. Inset: enlarged trace of a single AP.

To determine whether the non-adapting FS and adapting NFS phenotypes had the expected AP voltage properties, we plotted the AP waveforms and phase plots (dv/dt vs V) (Fig. 5B). As expected^55^, non-adapting FS cells had narrower AP waveforms, larger AHPs, and faster potential transitions, whereas NFS cells had broader and slower APs with less pronounced AHPs (Fig. 5C). Additionally, the two groups displayed expected differences in intrinsic electrophysiological properties, including input resistance, AP half-width, AP firing rate, pA at maximal AP firing rate, AP rise time, and AHP delay (Fig. 5C). Moreover, spontaneous APs, recorded at each cell’s resting membrane potential without stimulation, further confirmed the FS and NFS phenotypes (Fig. 5D) seen in the current injection experiments. Together, these results suggest that transplanted hMGE-pINs undergo prolonged maturation over months to years, developing the electrophysiological phenotypes consistent with non-adapting FS and adapting NFS INs.

### Transplanted hMGE-pINs form subclass-specific synaptic puncta

We next analyzed markers of synapse formation and connectivity associated with PVALB and SST interneurons (Supp. Fig. 6A) across the cortical layers. Endogenous SST-expressing interneurons typically target distal dendrites, while PVALB basket cells form perisomatic and proximal synapses on pyramidal cells and other interneurons (Supp. Fig. 6B)^20,100,101^. Using expansion microscopy (ExM)^102^, we analyzed the expression of human-specific TAU and/or SYP and observed that transplanted hMGE-pINs developed abundant synaptic puncta in layers I/II and IV/V at 7MPT (Supp. Fig. 6C-D). We hypothesized that layers I/II synapses were likely formed by human SST+ Martinotti cells, while layers IV/V signal corresponded to perisomatic synapses established by human PVALB-type basket cells (Supp. Fig. 6B). Consistently, in layers I/II, we identified inhibitory synapses (as shown by the close contact between human presynaptic SYP and postsynaptic Gephyrin), which were formed by SST+ hMGE-pINs targeting the distal dendrites of pyramidal cells (labeled by Pex5L, Supp. Fig. 6E-G). These axonal boutons expressed SST, SYNPR, and GRIN3A (Supp. Fig. 6G-I), which have been implicated in the specification and formation of synapses by SST+^103^. Furthermore, we detected axonal expression of the SST marker ARHGAP6 (Supp. Fig. 6J), a member of Rho GTPase activating protein family suggested to have an important role^104^.

In layer IV, we observed human axons surrounding the soma of endogenous neurons, with colocalized expression of markers suggestive of synapse formation, as shown by human SYP and the host postsynaptic marker GabaR1 (Supp. Fig. 6K-L). These soma-associated human synapses expressed LGI2, a protein involved in the formation of FS basket cell synapses^103^, as well as the potassium channels KIR3.1 (encoded by *KCNJ3*) and KV2.2 (encoded by *KCNC2*), both of which are enriched in the PVALB interneurons (Supp. Fig. 6A, M-O)^99,105^. These findings suggest that transplanted hMGE-pINs can form efferent synaptic connections with host neurons.

### Transcriptional evidence for diverse PVALB and SST subtypes, including SST interneurons with selective vulnerability during the early stage of Alzheimer’s disease

Pallial interneurons exhibit remarkable subtype diversity, conserved across adult cortical areas^10^. To assess the subtype diversity within hMGE-pIN grafts, we first analyzed the correlations between the snRNAseq clusters. Within the predicted PVALB subclass, cluster 9 is very distinct, whereas clusters 5 and 1 are highly correlated to each other, suggesting the presence of at least two PVALB populations (Fig. 6A). Cluster 7 is correlated with both PVALB and SST subclasses, consistent with overlapping marker expression patterns (Fig. 1J). Within the predicted SST subclass, there is a continuum including clusters 0, 4, 3, 2, and 8, suggestive of multiple related populations, whereas clusters 6 (NPY) and 10 (ADARB2) are transcriptionally distinct (Fig. 6A).

**Figure 6.**
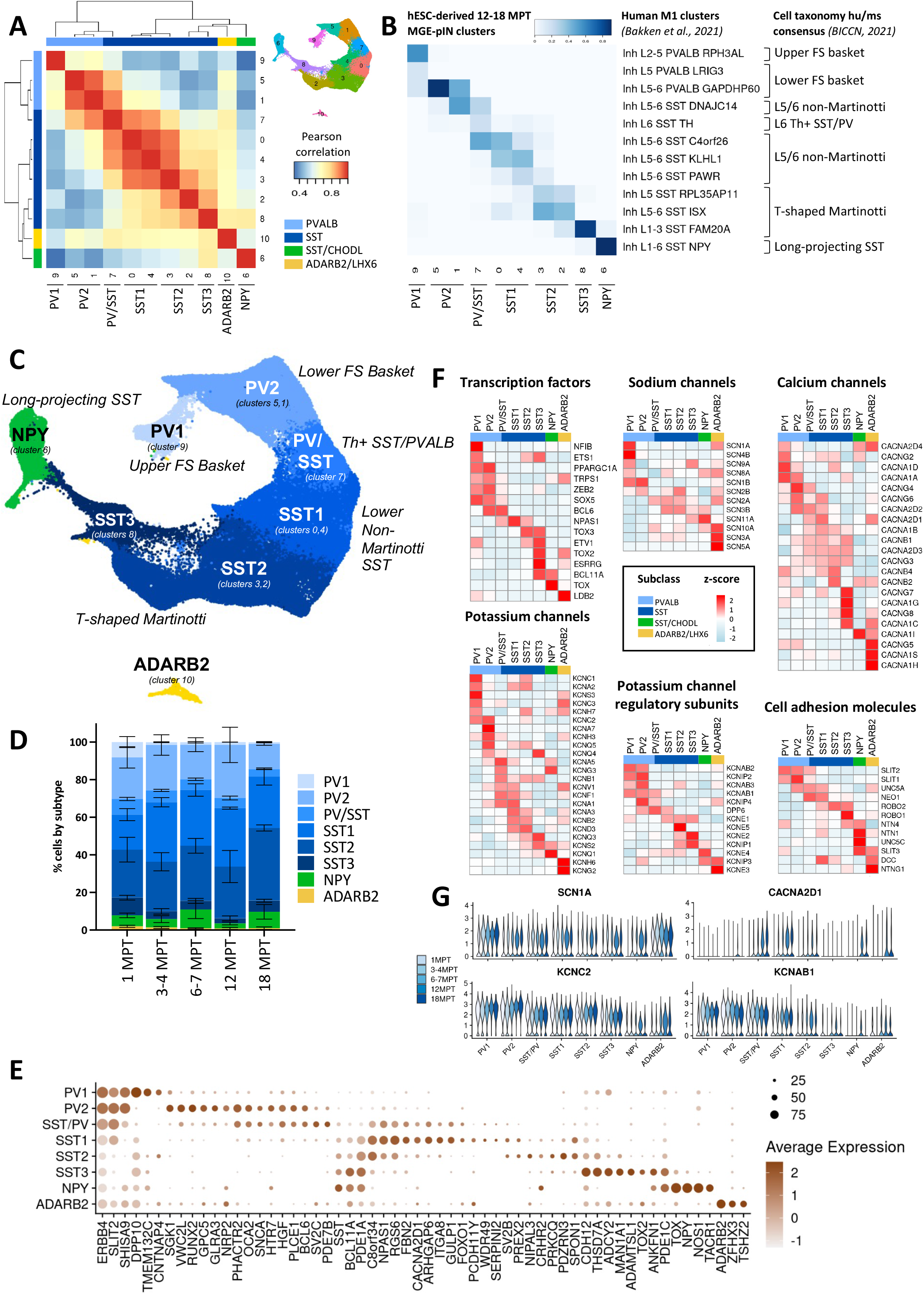
hMGE-pIN grafts comprise multiple PVALB and SST subtypes. (A) Pearson correlation of gene expression between clusters. (B) Distributions of predicted cell types for each 18 MPT cluster (columns) compared to endogenous adult interneuron transcriptional clusters (rows) defined in the cell census atlas of the mammalian primary motor cortex^9^. Rows include 10% or more correspondence between 18 MPT clusters and endogenous interneuron clusters. To the right, human cluster cellular taxonomy classification is shown, based on mouse/human cross-species analysis, where multimodal data (morphology, physiology and transcriptomics) were applied to define the mouse interneuron subtypes^9^. (C) Assignment of interneuron subtype identity. (D) Graft composition by subtype over time. Bar graphs shown mean ± SEM. (E) Expression of top subtype markers. (F) Expression of subtype-specific transcription factors, sodium, calcium and potassium voltage gated ion channels and regulatory subunits, as well as cell adhesion molecules involved in cell recognition^21^, in 12-18 MPT nuclei. (G) Expression of several voltage gated ion channels over time and across subtypes.

To investigate the subtype identities of the pallial interneuron clusters we mapped the oldest 12-18 MPT hMGE-pIN clusters in our study to the endogenous adult human cIN snRNAseq clusters defined in the M1 region of motor cortex^93^ (Fig. 6B; Table 5). The subpallial ADARB2 cluster (#10) was excluded from this analysis, since the corresponding endogenous populations are not represented in the cortical reference datasets. The identities of the reference clusters were defined by the human/mouse consensus taxonomy reflecting multimodal characterizations^9^. This analysis revealed that cluster 9 (PV1) matched human cluster “L2-5 PVALB RPH3AL”, which corresponds to the upper layer fast-spiking PVALB-type basket cells, whereas clusters 5 and part of 1 (PV2) matched human cluster “L5-6 PVALB GAPDHP60”, which corresponds to the deep layer fast-spiking PVALB-type basket cells (Fig. 6B, C). The superficial (PV1) and deep layer (PV2) PVALB subtypes could be distinguished by *NFIB* and *BCL6*, respectively (Fig. 6F), but shared other transcription factors (TFs), including *ETS1, PPARGC1A*, *ZEB2*, and *SOX5*, consistent with endogenous mouse and human cortical INs^21,93^.

Cluster 7 (PV/SST) shared transcriptional similarity with three human clusters, including two deep layer non-Martinotti SST clusters and L6 Th+ SST/PV populations (Fig. 6B, C), suggesting that this cell population may have the properties of both PVALB and SST subtypes, although it was grouped with the latter for quantification purposes (Fig. 1K-N). Clusters 0 and 4 (SST1) matched human clusters corresponding to deep layer non-Martinotti SST cells, while clusters 3, 2 (SST2) and 8 (SST3), matched the deep and superficial T-shaped Martinotti human clusters, respectively (Fig. 6B, C). All the SST interneuron subtypes expressed *RELN* (Fig. 1J), but could be distinguished by *NPAS1, NPAS3, ETV1,* and *BCL11A* TFs, respectively (Fig. 6F; Fig. 1O). The last SST population, cluster 6 (NPY), was distinguished by the *TOX* TF (Fig. 6F) and matched the human “L1-6 SST NPY” cluster, corresponding to MGE-derived long-range projecting (or LRP) neurons^80,9^ (Fig. 6B; Table 5).

Recent studies uncovered selective vulnerability of SST-type interneurons during the early phase of Alzheimer’s disease (AD) progression, prior to the onset of neuropathological or cognitive manifestations, including RELN-expressing populations which may confer resilience to cognitive decline^53,56,106^. Moreover, during the later phase of AD, characterized by an exponential increase in neuropathology and cognitive decline, there was a precipitous loss of some PVALB-type interneurons^106^. To determine whether any of these subtypes are present in the hESC-derived MGE-pINs grafts, we mapped transplanted interneuron clusters to the same foundational reference from the MTG as Gabitto et al.^106^. Many of the endogenous MGE-derived interneuron populations matched cells within specific clusters in the hMGE-pIN grafts (Supp. Fig. 7A, D), underscoring the potential for great transcriptomic diversity among *in vitro* derived interneurons. Most of the vulnerable SST subtypes depleted early in AD^106^ mapped to hMGE-pIN clusters 2 (SST2) and 8 (SST3), with human MTG clusters Sst-9, Sst-3, and Sst-11 being the most abundant (Supp. Fig. 7A-D). These *SST*+ *RELN*+ populations (Fig. 1J) correspond to Martinotti cells and constitute ∼20% of the human grafts (Fig. 6B- D). Of the PVALB subtypes depleted late in AD, human MTG clusters Pvalb-8 and Pvalb-14 mapped to hMGE- pIN cluster 9 (PV1) (Supp. Fig. 7A, C, D), corresponding to the upper layer FS basket cells (Fig. 6B, C). However, PV1 cells did not persist well beyond 1 MPT in the mouse xenotransplantation environment (Fig. 6D).

Collectively, these analyses demonstrated that hMGE-pIN grafts comprise diverse MGE subtypes, including upper (PV1) and lower (PV2) FS basket cells (∼21% total), SST/PVALB cells (∼7%), non-Martinotti cells (SST1, ∼32%), upper (SST3) and lower (SST2) Martinotti cells (∼33% total) and long-range projecting SST/NPY cells (∼6%), with the residual rare impurity being closely related MGE-derived GABAergic subpallial ADARB2/LHX6 cells (Fig. 6C, D). Importantly, none of the hMGE-pIN clusters matched any of the well-defined CGE-derived IN clusters (Table 5; Supp. Fig. 7D), highlighting specificity of the derivation method^72^.

### hMGE-pIN grafts comprise PVALB basket cells, as well as non-Martinotti, Martinotti and long-projecting SST cells, with corresponding histological features

To explore the molecular distinctions between the eight populations defined above, we analyzed expression of voltage-gated ion channels and cell adhesion molecules (CAMs) of the Netrin-Slit family (Fig. 6E-G) because these functional gene categories regulate synaptic communication and cell recognition. While voltage-gated ion channels in general are broadly expressed across cINs, the expression patterns of potassium (Kv), sodium (NaV) and calcium (CaV) voltage-gated ion channels and associated regulatory subunits are highly discriminative of specific subtypes^21^. Consistently, NaV genes (*SCN1A, SCN4B, SCN9A, SCN8A, SCN1B* and *SCN2B)* exhibited enrichment in PVALB basket cells (Fig. 6F) and developmental upregulation (Fig. 6G), similar to what has been described *in vivo*^21,107^. In addition, Kv channels and regulatory subunits exhibited subtype-specific enrichment, with notable examples associated with fast-spiking interneurons such as *KCNC1, KCNC2* (encoding Kv3), *KCNA2, KCNA7*, *KCNAB1, KCNAB2*, and *KCNAB3* (encoding Kv1-type potassium channel and modulatory subunits) (Fig. 6F, G). On the other hand, Martinotti cells, most clearly exemplified by SST3, showed distinct enrichment in several CaV genes, including *CACNG7, CACNA1G, CACNG8,* and *CACNA1C* (Fig. 6B, F). There was also subtype-specific expression of the Netrin-Slit family of CAMs (Fig. 6B), suggesting that this aspect of intercellular connectivity regulation is recapitulated in the context of hMGE-pIN xenotransplantation.

MGE-derived pIN subtypes can be further distinguished histologically by their morphologies, laminar distribution, and molecular markers (Fig. 7A). To distinguish the main PVALB, SST, and SST/NPY subclasses, we performed IHC for ERBB4/SST and NOS1/SST markers. At 7 MPT, we observed differential layer distribution, depending on marker expression patterns: SST-/ERBB4+ cells (consistent with PVALB subclass) preferentially populated layers IV and VI, SST+/NOS1- cells (consistent with SST interneuron subclasses) preferentially populated layers II-IV, whereas SST+/NOS1+ cells (consistent with SST/NPY subclass) preferentially populated deep layer VI (Fig. 7B), suggesting a notable degree of specificity in laminar distribution in the neocortex depending on cell fate, which is consistent with endogenous distribution patterns observed in mouse and human^4,8^.

**Figure 7.**
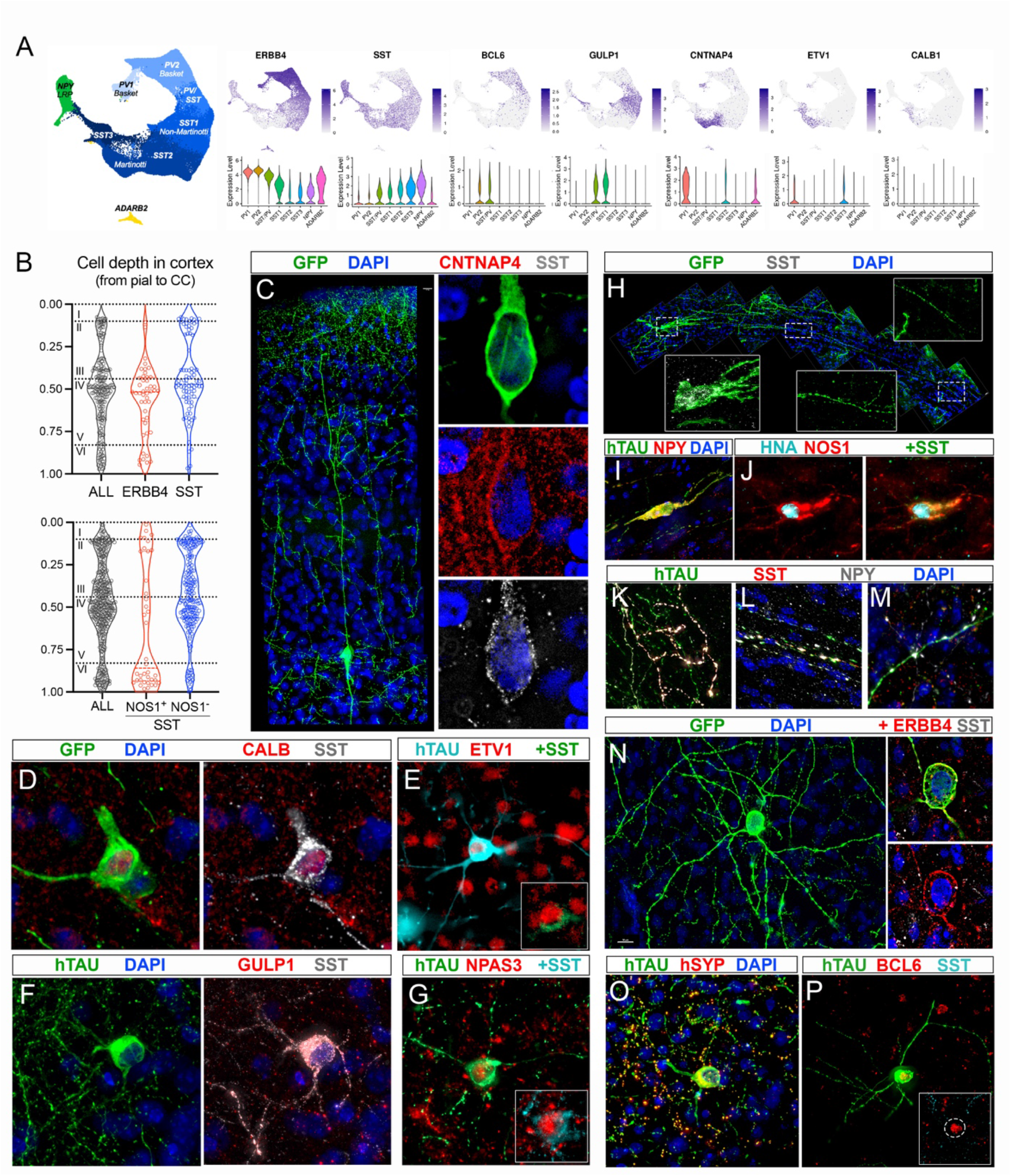
Grafted hMGE-pINs acquire histological features of known PVALB and SST subtypes. (A) Expression of markers used for histological characterization of human interneuron subtypes. (B-P) All images and quantifications were taken from animals at 7-15 MPT. Human cells were identified using either GFP or human-specific TAU antibody staining. (B) Relative positioning of hMGE-pINs in the cortex based on expression of ERBB4 / SST (upper panel, n=199 cells from 3 transplanted animals; 22.6% ERBB4+, 41.2% SST+) and SST / NOS1 (lower panel, n=396 cells from 3 transplanted animals; 49.4% SST+, 9.6% NOS1+). Dashed lines indicate the surface, layers I/II, III/IV, and V/VI transitions (from top to bottom). (C) Example of a GFP-labeled human neuron with Martinotti-like morphology – long ascending axon collaterals that reach and profusely ramify layer I. Insets showing SST and CNTNAP4 expression. (D, E) Human SST-positive cells co-expressing Martinotti markers CALB1 (D) and ETV1 (E). (F, G) Human SST-positive cells co-expressing predicted non-Martinotti subtype markers GULP1 (F) and NPAS3 (G). (H) Example of a GFP-labeled human neuron with long-range projection (LRP)-like morphology in layer VI with a long axonal collateral projecting parallel to the corpus callosum (inset shows SST expression in the soma and higher magnifications of the long projection). (I, J) Human cells with LRP characteristics frequently express SST, NPY, and NOS1. (K-M) Representative images of NPY+SST+ axonal projections, which are frequently found in the caudal hippocampus (medial DG-CA3 (K)), midline corpus callosum (L), and caudate putamen (next to the nucleus accumbens (M)). (N) Example of a ERBB4 positive, SST negative hIN with basket cell-like morphology in layer IV. (O) These cells form axonal arborizations (marked by human-specific SYP) around the somata of pyramidal neurons and interneurons. (P) Nuclear expression of the predicted basket subtype cluster marker BCL6. Inset shows BCL6 nuclear expression and lack of SST expression.

To characterize the morphologies of hMGE-pINs, cells expressing membrane-bound GFP were analyzed with expansion microscopy (ExM)^102^. Three types of morphologies in SST-expressing hMGE-pINs could be distinguished: Martinotti, non-Martinnotti, and LRP cells (Fig. 7C-M). Additionally, we identified basket-like morphologies in SST-negative cells (Fig. 7N-P). Martinotti-like human cells with pyramidal or oval shaped somas were characterized by bipolar or bitufted dendritic morphologies and long, ascending axonal collaterals that profusely ramified in layer I (Fig, 7C)^108,109^. These cells expressed low levels of SST together with CNTNAP4, CALB1, and ETV1 (Fig. 7A, C-E). Cells with non-Martinotti morphologies were found in layers II to V and were characterized by multipolar morphologies and co-expression of SST with GULP1 and NPAS3 (Fig. 7F, G)^110^. Cells with LRP morphologies were easily identified in layer VI and were further found dispersed in layers I to V (Fig. 7H). LRP cells were identifiable by their horizontal axonal trajectories running parallel to the corpus callosum, and by co-expression of SST with NOS1 and NPY (Fig. 7I-J)^80,111^. Furthermore, we observed NPY+/SST+ axon projections along the hippocampus, near the nucleus accumbens in the striatum, and traveling contralaterally through the corpus callosum in the forebrain midline (Fig. 7K-M). Conversely, ERBB4- expressing PVALB-type cells were negative for SST and displayed a multipolar dendritic morphology (Fig. 7N) with extensive human SYP-expressing axonal arborizations around nearby cells, reminiscent of basket cells (Fig. 7O)^108,112^. Although PV expression was found in a very small number of cells at late timepoints (> 17 MPT, data not shown), we observed co-expression of ERBB4 and BCL6 as early as 4 MPT (Fig. 7P), consistent with enrichment of these markers in the PVALB subclass (Fig. 1J; Fig. 7A). We did not observe any ERBB4-expressing cells with chandelier-like morphologies, consistent with snRNAseq results.

Taken all together, these results indicate that ESC-derived human MGE-pINs transplanted into mouse cerebral cortex display key characteristics of diverse endogenous human MGE interneurons, including transcriptional profile, morphology and layer-specific distribution, and provide strong evidence for subtype-specific integration into existing cortical circuits, along with progressive molecular and electrophysiological maturation.

## DISCUSSION

Our previous work laid the foundation for the current study by outlining the method for deriving highly pure GABAergic hMGE-pINs and describing their specification from NPCs, pre-transplant molecular identity, *in vitro* functional properties, and therapeutic effects in a rodent model of chronic focal epilepsy^72^. Building on this groundwork, we now present systematic analyses of hMGE-pINs from 1 to 18 months after transplantation using snRNAseq, long-term electrophysiological recordings, and histological characterization. This study represents a comprehensive, quantitative *in vivo* characterization of a human PSC-derived neuronal cell therapy candidate under clinical evaluation for the treatment of drug-resistant mesial TLE (NCT05135091). We demonstrate that after transplantation, hMGE-pINs exhibit high on-target purity, undergo intrinsic and reproducible differentiation trajectories, and maintain stable composition made up of authentic pallial MGE interneurons. Our approach differs from other ongoing cell therapy clinical trials for neurological disorders like ALS, multiple sclerosis, Parkinson’s, Huntington’s, Alzheimer’s, and spinal cord injury, where multipotent stem cells, such as mesenchymal or neural stem cells, or mixed populations of regionally patterned progenitors and neurons are transplanted without precise control over their proliferation and differentiation potential^113–117^. By transplanting correctly specified and committed postmitotic GABAergic MGE-type pallial interneurons, we demonstrate reproducible and durable control over graft composition, setting a new standard for targeted cellular therapies.

Our findings suggest how different aspects of interneuron functionality may contribute to the modulation of hyperexcitable circuits. While MGE-pINs secrete GABA at the time of transplantation, it takes 5-7 months for significant seizure suppression in the murine model of focal TLE^72^, indicating that tonic GABA secretion alone is not sufficient and that integration with the host circuitry may be critical for maximum efficacy in the xenotransplantation context. The kinetics of transcriptional upregulation of synaptic transmission machinery uncovered in this study could explain the observed delay in functional recovery. Interestingly, at the time that hMGE-pINs are efficacious against kainic acid-induced epilepsy (5-7 MPT), most of the grafted human interneurons exhibit immature electrophysiological properties and lack robust *SST* or *PVALB* expression, indicating that fully developed electrochemical and physiological properties are not required for the therapeutic benefit in this model.

Despite great efforts over the past decade, generation of PVALB-type interneurons *in vitro* from human hPSCs has been challenging, with rare observations of PV-positive cells and no convincing electrophysiological, morphological or single cell transcriptomic evidence published thus far ^40,65–74,87,88,90,118–124^. In our previous study, we reported rare instances of PV-positive cell fate post-transplantation and expression of perineuronal nets associated with fast-spiking neurons^72^, but a large proportion of the grafted GABAergic LHX6-expressing cells remained undefined in terms of subclass fate. Here, using snRNAseq we demonstrate that all grafted human cells are MGE-derived GABAergic neurons, with over 97% of them having on-target pallial SST or PVALB identity across the lifespan of the xenotransplantation mouse model. Comparative transcriptomic and other supporting modalities further reveal that a substantial proportion of grafted cells (∼20%) have reproducible and stable PVALB identity that matches human basket cells, even though *PVALB* mRNA itself is not yet expressed at these relatively early stages of human interneuron development. In fact, the lack of *PVALB*/PV expression in immature human PVALB-type pINs should be expected, given the protracted developmental timeline of this late-born population, with robust expression levels reported only after ∼10 years of age in humans, and the documented influence of sensory input on their maturation^36,94,97,125–127^. Moreover, it is possible that the rodent environment somehow inhibits *PVALB*/PV expression in human interneurons, consistent with recent studies showing that mouse MGE interneuron progenitors only expressed PV when grafted onto human organotypic slices or 3D organoids, but not the mouse counterparts^128^.

In this study, we relied on multiple lines of mutually supportive evidence, including comparative single cell transcriptomics, electrophysiology, morphology and expression of known markers, to identify PVALB and SST subclasses generated from hESCs. Most of the major MGE-derived interneuron populations were recapitulated in our grafts, with consistent composition across lots and timepoints post-transplant, including Martinotti and non-Martinotti SST cells, long-range projecting SST/CHODL (NPY) cells, and PVALB basket cells. The only notable exceptions were the LAMP5-LHX6 neurogliaform (NGC) and chandelier cells, which to our knowledge have not yet been reported in any hPSC-derived models. However, the exact number of subtypes could not be determined with certainty, presumably due to the relatively young age of human interneurons at 18 MPT and the lack of extrinsic factors that may contribute to human-specific interneuron maturation and transcriptional refinement^97,98,129^. Despite this limitation of the rodent host environment, many essential genetic programs responsible for cell identity were recapitulated in the xenotransplantation model. Moreover, we observed a gradual upregulation of vesicular transport, membrane potential regulation, and synapse formation modules, reaching expression levels consistent with human postnatal developmental stages by the end of our study.

Our slice recordings indicate that hMGE-pINs mature into non-adapting fast-spiking and adapting non-fast-spiking interneurons over months to years, exhibiting expected action potential (AP) and intrinsic properties, such as firing rates, adaptation, waveforms, kinetics, afterhyperpolarization (AHP) amplitudes, and input resistance. The electrophysiological properties of our hMGE-pINs closely resemble endogenous human MGE interneuron cells^130^, with some cells displaying firing rates ranging from 100–200 Hz and both adapting and non-adapting AP phenotypes.

Unexpectedly, we observed distinct interneuron subclasses rapidly emerging from the homogeneous population of postmitotic interneurons within 2-4 weeks after transplantation, well before full neurochemical and functional maturation. While 30 DIV co-culture with mouse astrocytes led to a limited subclass differentiation, 14 days of xenotransplantation facilitated rapid subclass specification, suggesting that interactions with the host neuronal and glial populations and/or access to brain neurotrophic factors^131^ may activate intrinsic programs in the immature postmitotic interneurons. Thus, the xenotransplantation approach combined with comparative transcriptomics, provide a powerful platform for studying the mechanisms of human pIN subtype specification, maturation and function, particularly in the context of neurodevelopmental disorders associated with pIN abnormalities, such as epilepsy, autism, schizophrenia, and ADHD^132^.

The ability to identify cell fate within weeks after *in vivo* transplantation overcomes the limitations posed by the slow electrochemical maturation of human interneurons. This approach not only facilitates the efficient development of regenerative cell therapies for neurological diseases but also may serve as a platform for future mechanism-of-action and disease-modeling investigations. As single-cell profiling datasets from various neurological disorders become increasingly available, one can envision tailoring cell therapies to fit specific target brain regions or structures based on the endogenous cell types affected by the disease. For instance, recent evidence points to specific vulnerability of several SST interneuron subtypes, including ones represented in our hMGE-pINs, in the subgranular layer of the middle temporal gyrus during the early AD phase, with the loss of these interneurons potentially accelerating cognitive decline^56,106^. These observations point to a possible SST-enriched hMGE-pIN cell therapeutic approach that merits further exploration. In the case of TLE, the precise relationship between the human interneuron subtype composition in the grafts and disease-modifying effects remains unclear. The derivation protocol used for hMGE-pINs in this study primarily generates SST subtypes, comprising ∼70% of the population, while the rest is split between PVALB-type basket cells and long-range projecting (LRP) NPY cells. However, other subtypes such as chandelier or NGC cells, that could play an important role in certain forms of epilepsy and other neurological disorders, have yet to be generated *in vitro* from hPSCs. Future studies may seek to determine which subtypes are critical for seizure suppression in epilepsy, as well as the development of new differentiation protocols to tailor specific interneuron subtypes for improving other disease phenotypes. While this will be a challenging endeavor, the necessary tools and techniques are now in place to make significant progress in these areas.

## ACKNOWLEDGEMENTS

This work was supported by the California Institute for Regenerative Medicine (CIRM; grants: DISC2P-11700, DISC-10525 and TRAN-11611, principal investigator: CRN). We thank Neurona co-founders John Rubenstein, Arnold Kriegstein, Arturo Alvarez-Buylla for guidance and review of the manuscript, as well as Thomas Faust for helpful feedback.

## AUTHOR CONTRIBUTIONS

Conceptualization: MB, RZ, LF, YM, CRN; Methodology (direct): MB, RZ, LF, CC, GS, DC, JS; Methodology (indirect): MB, GS, MS, YM, SH, SK, WA, SL, MW, AN, DA, BF; Software: RZ, JS; Data Analysis: MB, RZ, LF, CC, GS, DC, JS; Investigation: MB, LF, CC, GS, DC, JS, MS, WA, YQ, BF, OK, MG, VH, JHJ, TK; Writing original draft: MB with contributions from LF, RZ, CC, GS, and JP; Editing: MB, RZ, LF, CC, GS, DC, JS, MS, YM, SH, MW, TK, AB, CP, JP, CRN; Visualization: MB, RZ, LF, CC, JS; Supervision: MB, LF, GB, CP, JP, CRN. All authors reviewed and approved the manuscript.

## STAR METHODS RESOURCE AVAILABILITY

### Lead contacts

Further information and requests for resources should be directed to and will be fulfilled by the lead contacts, Marina Bershteyn or Cory Nicholas.

### Materials availability

This study did not generate new unique reagents.

### Data and code availability

Single-cell RNA-seq data (original and reanalyzed) will be deposited in NCBI’s Gene Expression Omnibus (GEO) and will be publicly available as of the date of publication (*future GEO Series accession number*). All original code will be deposited and publicly available as of the date of publication. DOIs will be listed in the key resources table. To review GEO accession: go to- https://www.ncbi.nlm.nih.gov/geo/query/acc.cgi?acc<u>=</u>*** Enter token- (TBD). Any additional information required to reanalyze the data reported in this paper is available from the lead contact upon request.

## EXPERIMENTAL MODEL AND STUDY PARTICIPANT DETAILS

### Institutional oversight

All activities, procedures and materials used were reviewed and approved by Neurona’s Stem Cell Research Oversight Committee and Institutional Animal Care and Use Committee (IACUC).

### hESC lines

A proprietary male cGMP-grade human embryonic stem cell (hESC) line that is listed in the NIH hESC Registry as an “Approved Line” (eligible for use in NIH-supported research) was used to generate research-grade working cell banks (rWCBs). Donor consent was obtained for use of the cells in research, clinical and commercial development. Donor screening was performed using FDA’s Donor Eligibility Guidelines for HCT/Ps. Karyotypic analysis and pluripotency marker expression (TRA-1-60, OCT-4, and NANOG) were assessed across all hESC cell banks, demonstrating genomic stability and high purity. Pluripotency was demonstrated by *in vitro* differentiation of the cells to endoderm, ectoderm, and mesoderm lineage; and the demonstration of teratoma formation when undifferentiated hESCs were engrafted *in vivo*. The pluripotent stem cell banks were tested for sterility and found negative for relevant viruses, mycoplasma, and adventitious agents. The hESC WCBs were tested for genetic stability per ICH Q5D (ICH_Q5D 1998)^72^. The cell line was adapted to a feeder-free, xeno-free culture system using human recombinant vitronectin coated dishes, Essential 8 medium kit and dissociation reagent ReLeSR, with daily media changes.

### Animals

Activities described in this study were performed in accordance with the active Neurona Therapeutics IACUC protocol covering temporal lobe epilepsy and adhered to the ARRIVE guidelines. Neonatal (P1-P2) Scid-beige (*CB17.Cg-PrkdcscidLystbg-J/Crl*) mice were obtained from Charles River Labs. After weaning, mice were assigned a unique identifier number which was marked on the mouse tail. Animals were maintained in the Neurona Therapeutics Animal Facility under standard temperature and humidity on a 12:12 light/dark cycle. Bottled water and standard chow were provided ad libitum. Colony health was monitored through monthly diagnostic sample submission for microbiology.

## METHOD DETAILS

### Generation of MGE pallial-type interneurons from hESCs

Cells were differentiated as described previously^72^. Briefly, rWCBs of hESCs were thawed (one vial of rWCB for each differentiation experiment) and expanded for 5 days as described above. To initiate patterning into definitive neuroectoderm, hESCs were dissociated with StemPro Accutase and resuspended in differentiation media (50% Neurobasal-A; 50% DMEM/F12; 1X GlutaMAX; 1X B-27 without Vitamin A; 1X N2 supplement-B; 1X Penicillin-Streptomycin; 200 μM L-Ascorbic Acid; 55M β-Mercaptoethanol; 1/2X MEM non-essential amino acids) supplemented with Rho kinase inhibitor Y27623 (10 μM), and dual SMAD inhibitors SB431542 (10 μM) and LDN193189 (250 nM). Forebrain identity was promoted with WNT pathway inhibition using XAV939 (10 μM), and ventral forebrain MGE progenitors were induced with SHH pathway activation using SAG (100nM). After two weeks of patterning, MGE progenitors, defined by co-expression of SOX1, FOXG1, NKX2-1^72^ were expanded and grown for a total of four more weeks. MGE neuronal differentiation was indicated by upregulation of postmitotic markers LHX6 and LHX8, with the MGE pallial lineage (MGE-pIN) being characterized by downregulation of LHX8 and NKX2-1 and co-expression of LHX6 with MAF, MAFB, ERBB4, CXCR4, and CXCR7^72^. During the last weeks of differentiation, MEK pathway inhibitor PD0325901 (2 μM) as well as inhibitors of CDK (PD0332991, 2 μM) and NOTCH pathways (DAPT, 10 μM), though not required for MGE-pIN induction, were implemented to promote downregulation of subpallial MGE marker LHX8 and cell cycle exit, respectively^72^. At the end of the differentiation process, cells were harvested and dissociated to single cells using TrypLE™ Select Enzyme and Benzonase (1/10,000). ERBB4-positive cells were sorted to enrich MGE-pIN purity with a biotinylated primary antibody against human ERBB4 and magnetic sorting using the CliniMACS. Cells were cryopreserved using the CryoMed controlled-rate freezer before further storage in vapor-phase liquid nitrogen. All analytical assays and experiments were performed after thawing sorted hMGE- pIN cell lots from cryopreservation. The experiments described herein used non-GMP-grade hESC rWCBs to produce MGE-pIN cell lots, and do not represent the clinical candidate (NRTX-1001) that is being evaluated in a clinical trial for drug-resistant epilepsy. All the relevant materials are listed in the key resource table. Further details are described in Bershteyn et al.^72^

### Cell Transplantation

Neonatal (P1-P2) Scid-beige (*CB17.Cg-PrkdcscidLystbg-J/Crl*) mouse pups were anesthetized via hypothermia until the pedal reflex disappeared and then placed on a stereotaxic platform. Dissociated hESC-derived MGE-pINs were delivered using a pulled glass needle (70-90 μm diameter) connected to a microinjection system (Narishige). For RNAseq experiments, pups were injected in 3 sites per hemisphere (AP: 2.5 mm / ML: +/-1.1 mm; AP: 1.7 mm / ML: +/-1.0 mm; AP: 0.9 mm / ML: +/-0.8 mm, with respect to lambda) at a dose of 75E3 cells per site. For histological experiments, pups were injected at a single site per hemisphere (AP: 1.7mm / ML: +/-1.0mm, with respect to lambda) at a dose of 50E3 cells per site. All injections were made at 0.3-0.5 mm deep from the skull. After injection, recipients were placed on a heating pad until warm and active, at which time they were returned to their dams until weaning (P21).

### Tissue Dissection

Animals were fully anesthetized with 2% isoflurane, decapitated, and the brain was removed into ice-cold high-sucrose brain slice cutting solution (HS-BSCS: 75 mM sucrose, 2.5 mM KCl, 4 mM MgCl2, 1.25 mM NaH2PO4, 24 mM NaHCO3, 25 mM glucose, 0.5 mM CaCl2, 85 mM NaCl oxygenated with 95% O2 / 5% CO2 to pH of 7.4). Cortical tissue was then dissected in cold oxygenated HS-BSCS, chopped into small pieces (< 1 mm^3^), flash-frozen, and stored at -80°C before processing for dissociation.

### Nuclei isolation and fluorescence-activated nuclei sorting (FANS)

Dissected tissue was retrieved from -80°C storage, placed on dry ice and weighed to determine the combined total mass per sample. Homogenization buffer (HB) was prepared fresh for each experiment using reagents from the Nuclei Isolation Kit (Nuclei PURE lysis buffer + 0.1% Triton X-100), supplemented with DTT (1mM ) and Protease Inhibitor cocktail (1X), and kept on ice. HB volume was prepared at 25 mL per 125-150 mg brain tissue. Tissue was resuspended with cold HB and transferred using sterile transfer pipettes into a pre-chilled dounce homogenizer. Dissociation was done with ∼20 strokes, keeping the homogenizer in ice. Lysates were diluted to the total volume of HB based on original sample mass and incubated on ice for 20-30 minutes prior to being filtered through a 40 μm strainer into 50 mL conical tubes (up to 50 mL lysate volume per tube) to remove large debris. Lysates were centrifuged at 900G for 5 minutes at 4-15°C. The supernatants were discarded, and each pellet containing nuclei was resuspended with 1 mL of cold Nuclei Wash Buffer (NWB): D- PBS supplemented with Ultrapure bovine serum albumin (1%) and RNAse Inhibitor (0.2 U/μL), triturated 10 times and filtered into FACS tubes through the strainer in the cap. Pellets from the same sample were combined in one FACS tube, and another 1 mL of NWB was used to rinse the original 50 mL conical tubes before being added to the rest of the sample. Samples containing nuclei were stored in ice for 20 minutes, during which time nuclei count was performed using NC-200.

To label human nuclei, primary mouse anti-human STEM101 antibody was added to the samples (1:500) and incubated for 1 hour on ice. Note, the staining reaction was validated in the range of 3E6-10E6 nuclei/mL, and typically did not require concentration adjustments from the initial dilution in the NWB. Nuclei counts were recorded to monitor yields throughout the process and across experiments. Following primary antibody incubation, samples were centrifuged at 900G for 5 minutes at 4-15°C. Nuclei pellets were resuspended again with NWB (0.5-1X volume compared to the original NWB volume used for primary antibody incubation). Secondary donkey anti-mouse AF647 antibody was added (1:1000) and incubated for 30 minutes on ice in the dark. Following secondary antibody incubation, samples were topped off with NWB, centrifuged at 900G for 5 minutes at 4-15°C, resuspended with NWB (0.5-1X volume compared to the original NWB volume used for primary antibody incubation) and filtered into FACS tubes through the strainer in the cap. Sytox Blue Dead Cell Stain was added (1:1000) and samples were enriched for STEM101-AF647+ nuclei using the BD FACSAria cell sorter. Samples were first gated to exclude debris and select for nuclei (FSC-A vs BV421-A, left panel below), followed by gating to select for single nuclei (FSC-A vs SSC-A, middle panel below), finally the single nuclei were gated to select for and sort the STEM101-AF647+ population (APC-A vs FSC-A, right panel below).

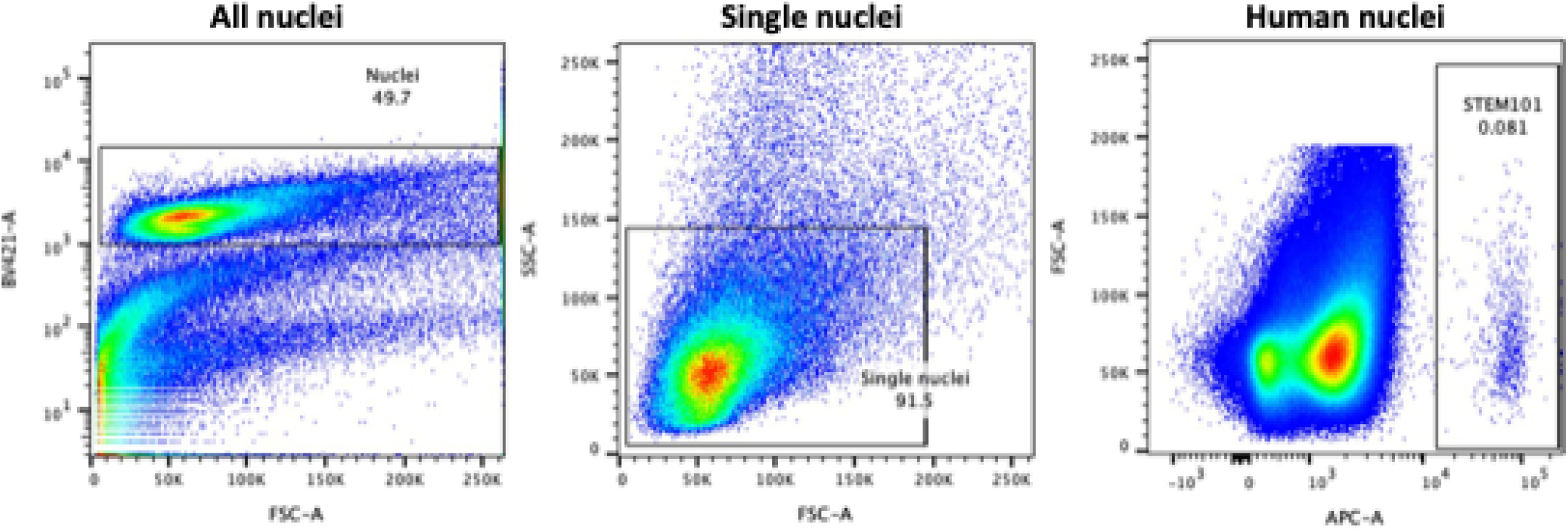

### Single cell/nuclei RNA sequencing (sc/snRNAseq)

To capture single nuclei for RNA sequencing, the entire sample that was recovered from the sort was loaded onto 10X Genomics microfluidic chips. Typical concentration after sort was 100-200 nuclei/μL, and the entire volume was split across multiple lanes, per manufacturer instructions.

To capture single cells for RNA sequencing, samples were thawed on the day of capture, viability was measured using NC-200 (>90% post-thaw viability was routinely achieved and expected). Samples were diluted to 1E3 cells/μL in culture medium. To ensure high quality single cell data, High Sensitivity RNA assay was performed, and ambient RNA was measured using Qubit 3.0 Fluorometer, with the cut-off of <3 ng/μL ambient RNA. Samples were loaded per manufacturer instructions, for target recovery of 1E4 cells/sample.

Individual cells/nuclei were barcoded using the chromium controller, cDNA and sequencing libraries were prepared using next GEM 3’ V3.1 kits (10x Genomics). Quality control and cDNA/library quantification was done using the Agilent Tapestation 4200. Libraries were shipped overnight on dry ice to SeqMatic, LLC (Fremont, CA) for pooling and sequencing. Pooled libraries were quantified using Lightcycler 96 qPCR and sequenced by the Illumina NovaSeq V1.5 or NovaSeq X Plus, aiming for 15-20E3 reads per cell sequencing coverage. Typically, ∼8E3-1.2E4 cells were captured per sample.

### RNA fluorescence in situ hybridization (FISH)

RNA FISH was performed on fixed, frozen mouse brain sections using RNAscope Multiplex Fluorescent Reagent Kit v2 Assay according to the manufacturer’s instructions. Briefly, 40 µm sections were post-fixed with 4% PFA and dehydrated with 50%, 70% and 100% ethanol. Sections were pretreated with hydrogen peroxide and target retrieval was performed. Following a 30-minute protease treatment, sections were hybridized to probes against human targets: GAD1, NPY, SLIT2, SST, and MALAT1. MALAT1 probe was cross validated not to recognize the mouse MALAT1 gene. Further amplification was performed according to the assay kit protocol, and Opal 520 (Akoya Biosciences cat# NC1601877), Opal 570 (Akoya Biosciences cat# NC1601878) and Opal 690 (Akoya Biosciences cat# NC1605064) dyes were used to fluorescently label the probes. Tile scan images were acquired using the Zeiss AxioScan slide scanner microscope. Analysis was performed manually using the Zeiss Zen microscopy software.

### Extended culture on glial feeders (30 DIV) for snRNAseq

For the extended in vitro culture experiment with sequencing, primary mouse glial feeder cells were isolated from E14.5 CB17 mouse embryos. Mouse cortices (6-12) were dissected, cut into pieces using sharp blades and placed in a 15 mL tube containing Leibovitz L-15 medium with DNase (1%). Once all cortices were collected, they were enzymatically dissociated with 0.25% Trypsin-EDTA/DNase, quenched with 20% Fetal Bovine Serum (FBS) and centrifuged at 350G for 5 minutes. The pellet was resuspended in plating media (Neurobasal-A medium supplemented with 1X B-27 without Vitamin-A, 1X GlutaMAX, 1X Penicillin-Streptomycin, 200 µM Ascorbic Acid) with 2% FBS and 1% DNase, mechanically dissociated and passed through a 40 μm strainer. Cells were counted and seeded at 0.2E6 cells/cm^2^ into cell culture vessels pre-coated with Poly-L-Ornithine, Laminin and Fibronectin (PO/Fib/Lam). Cells were maintained in culture over five days with media changes every other day using plating media (described above) supplemented with 20 ng/mL brain derived neurotrophic factor (BDNF) and 2% FBS. Once cultures were approximately 85% confluent, they were treated with 2 pulses of 5nM Ara-C over 7 days to inhibit proliferation.

hMGE-pINs were plated on Ara-C treated primary mouse glial feeder cells and maintained in culture with 3 media changes per week using plating media (described above) supplemented with 20 ng/mL BDNF. After 28 days of co-culture, cells were harvested and dissociated into single cells using 0.25% Trypsin-EDTA. Nuclei were isolated and human nuclei were sorted and captured for snRNAseq, as described above.

### In vitro calcium imaging

hMGE-pINs and primary mouse astrocytes (ScienceCell, NC1175467) were seeded in a 384-well plate at a neuron-to-astrocyte ratio of 5:1, at a total cell concentration of 36,000 cells per well. Before seeding the cells, plates were coated with PO/Fib/Lam. Neurons were transduced on day of seeding with Synapsin-GCaMP8m lentiviral construct purchased from VectorBuilder. Co-cultures were fed every other day with cell culture media as follows: BrainPhys, Neurocult SM1 supplement (1X), L-ascorbic acid (200 μM), N-2 supplement (1X), Antibiotic-Antimycotic (1X), laminin (1 μg/mL), BDNF (20 ng/mL), glial derived neurotrophic factor (GDNF, 20 ng/mL), cyclic AMP (cAMP, 500μM), and FBS (0.5%).

Imaging was done using the ImageXpress Micro Confocal instrument from Molecular Devices. On the day of imaging, the media was replaced with BrainPhys Imaging (1X) with the same components as listed above. A baseline recording was taken for 30 seconds, followed by compound addition after removing the plate from the instrument, which was then equilibrated for 20 minutes back in the instrument before a final 30-second post-compound recording. The images were then taken offline to be processed using a custom calcium imaging pipeline. The open source Suite2P package was used to identify and extract calcium traces, and the Cascade package was used to predict firing rate^133,134^.

### Histology

Brain tissue was sectioned at 40 µm on a slide microtome and processed for immunohistochemistry. Sections were first treated for antigen retrieval (5–10 min with boiling 0.01 M citrate buffer (pH = 6)), followed by a blocking step (one hour with 5% goat serum and 0.1% Triton X-100 in D-PBS), and incubated with primary antibodies overnight at 4°C, diluted in blocking buffer. After washing three times in D-PBS + 0.1% Triton X-100, sections were incubated with secondary antibodies (2 hours in blocking buffer), washed four times in D-PBS + Triton X-100, and mounted onto glass slides. Primary and secondary antibodies are listed in the key resource table.

Expansion microscopy was performed as described in Asano et al., 2020^102^. Briefly, brain sections were incubated in a Acryloyl-X, SE solution (overnight, 0.1 mg/mL in D-PBS at room temperature). After two washes with D-PBS, a monomer solution (1× D-PBS, 2 M NaCl, 8.625% (w/w) sodium acrylate, 2.5% (w/w) acrylamide, 0.15% (w/w) N,N′-methylenebisacrylamide, 0.01% (w/w) 4-hydroxy-2,2,6,6- tetramethylpiperidin- 1-oxyl (4- hydroxy-TEMPO), 0.2% (w/w) ammonium persulfate, and 0.2% (w/w)t etramethyl-ethylenediamine (TEMED) accelerator) was used to gelate brain sections before expansion (2-3 hours in a humidified 37°C incubator). For expansion, gels were treated with digestion solution overnight (Proteinase K in 50 mM Tris (pH 8), 1 mM EDTA, 0.5% Triton X-100, 1 M NaCl) at room temperature. Digested gels were then placed 3-4 times in deionized water for 2 hours at room temperature and mounted for imaging using a Leica DMi8 microscope with a 10 and 20X objective.

### Acute brain slice preparation

Electrophysiological experiments were conducted on acutely prepared brain slices containing cortical regions from 12 MPT to 20 MPT mice. Mice were anesthetized with avertin (tribromoethanol, 0.15 ml per 10 g body weight) and perfused with ice-cold dissection solution containing (in mM): 93 N-Methyl-D-glucamine (cat#M2004, Sigma-aldrich), 2.5 KCl (cat#P3911, Sigma-aldrich), 1.2 NaH_2_PO_4_ (cat#S0751, Sigma-aldrich), 30 NaHCO_3_ (cat#S6014, Sigma-aldrich), 20 HEPES (cat#H4034, Sigma-aldrich), 25 D-(+)-Glucose (cat#G8270, Sigma-aldrich), 5 Ascorbic acid (cat#1043003, Sigma-aldrich), 2 Thiourea (cat#T7875, Sigma-aldrich), 3 Sodium Pyruvate (cat#P2256, Sigma-aldrich), 12 N-acetyl-L-Cysteine (cat#A9165, Sigma-aldrich), 10 MgSO_4_(cat#M2643, Sigma-aldrich), and 0.5 CaCl_2_ (cat#C5670, Sigma-aldrich) with a pH of 7.2-7.4. The brain was then quickly removed from the skull and cooled in the ice-cold dissection solution. 300 μm coronal slices containing the cortex were sectioned and the slices were bisected down the midline and transferred to an incubation chamber containing the dissection solution warmed to 35°C. The sodium concentration in the chamber was increased gradually depending on the age of the mice. The method was described in Ting et al, 2018^135^. After the sodium concentration was increased to the same level of the recovery artificial cerebrospinal fluid (ACSF) containing (in mM): 92 NaCl (cat#S9625, Sigm-aldrich), 2.5 KCl, 1.2 NaH_2_PO_4_, 30 NaHCO_3_, 20 HEPES, 25 Glucose, 5 Ascorbic acid, 2 Thiourea, 3 Sodium Pyruvate, 12 N-acetyl-L-Cysteine, 2 MgSO_4_, and 2 CaCl_2_, the slices were transferred to the recovery chamber containing the oxygenated ACSF at room temperature for at least 1 h to allow for recovery prior to beginning recording sessions.

### Whole-cell patch clamp recordings

Hemi-slices were transferred one-at-a-time from the recovery chamber to the recording chamber, where they were submerged and perfused with ACSF at a rate of ∼2 ml/min. Transplanted channelrhodopsin or eGFP-expressing interneurons in the cortex were located under fluorescence illumination and subsequently targeted for whole-cell patch clamp recordings. Patch pipettes with a resistance of 3-6 MΩ were fabricated from borosilicate glass (ID 1.2 mm, OD 0.86 mm; Garner Glass) on a horizontal puller (P-97, Sutter Instrument) and filled with an internal patch solution containing (in mM): 130 potassium D-gluconate (cat#G4500, sigma-aldrich), 10 KCl, 1 NaCl, 1 MgCl_2_, 1 CaCl_2_, 10 EGTA (cat#E4378, Sigma-aldrich), 10 HEPES, 2 Mg-ATP (cat#A9178, Sigma-aldrich), 0.5 Na-GTP (ca#51120, Sigma-aldrich); the pH was adjusted to 7.3 with KOH and the osmolarity was adjusted to 300 mOsm with D-sorbitol. Whole-cell patch clamp recordings were obtained with a Multiclamp 700B amplifier (Molecular Devices), digitized at 10 kHz. Data analysis was done with pClamp 10 (Axon Instruments). Resting membrane potential was determined upon initiating whole-cell configuration. The action potential waveform kinetics were analyzed using the first action potential occurring at the rheobases. In the photo stimulation test, we used continuous, 10 Hz, 20 Hz, 40 Hz and 80 Hz LED stimuli (intensity around 1mW) to trigger cell spiking. The maximum AP frequency stimulated by continuous stimuli was calculated by intervals between every two APs. For interneuron subtype characterization, we group those neuron firing frequencies keep increasing in response to a series of current injections into the non-adapting FS group and those neurons show firing frequency adaptation into the adapting NFS group. The postsynaptic currents were recorded at –70 mV.

## QUANTIFICATION AND STATISTICAL ANALYSIS

### scRNAseq analysis for pre-transplant cells

Cell Ranger v4.0.0 (10x Genomics) was used to demultiplex FASTQ files for each sample, align reads to the human GRCh38 genome downloaded from 10x Genomics, and quantify the expression levels for each gene in each cell from each sample. R version 4.0.3 and Seurat v4.3.0 were used to quality control, normalize, cluster, and visualize the cells from all samples. Cells were filtered for quality by having at least 1000 expressed genes, at least 1000 unique reads, no more than 20% reads of mitochondrial genes and no more than 40% reads of ribosomal genes. The data were normalized by the ‘LogNormalize’ method and the mean expression of genes between cells were scaled to 0. The top 3000 genes were identified by ‘FindVariableFeatures’ with ‘vst’ selection method for downstream integration analysis. Seurat Canonical Correlation Analysis (CCA) integration workflow was applied to remove the batch effects related to cell capture and library-making procedures across cell lots. Principal component analysis (PCA) was applied, 30 principal components (PCs) were selected, and cells were clustered with Seurat standard workflow. Cells were visualized in a 2- dimensional space of UMAP (Uniform Manifold Approximation and Projection), and colored by groups or cell types, where each dot represents one cell. Groups of cells with similar gene expression form a cluster. Top markers for each cluster were identified by using the Seurat ’FindAllMarkers’ method with Wilcox test. The cell type of each cluster was determined by the expression of typical cell markers together with the predictions with the human and chimpanzee developing brain scRNAseq data. Gene expression was visualized with violin-plot, dot-plot and feature-plot built in Seurat package.

### Characterization of pre-transplant cell types by snRNAseq data from developing brain

To compare our hESC-derived interneurons with human developing GE, published snRNAseq datasets^35^ were downloaded from GEO dbs, and processed to identify MGE, CGE, LGE and other clusters using Seurat, as described in the original report. PCA was applied and the reference UMAP models of cell types and human developmental ages were built, and the cell types and the pseudo-developmental ages of the hESC-derived interneurons were predicted by following the Seurat reference mapping procedure^86^. Seurat identified anchor correspondences between the query datasets and the reference, annotated query cells by using a weighted vote classifier based on the reference cell identities. Multiple anchors were considered for the classification of each query cell and the predictions were informed by a cell’s local neighborhood. The anchor scoring approach of Seurat provides a quantitative score for each cell’s predicted label. Cells classified with high confidence will receive consistent votes across anchors, whereas cells with low confidence, including cells not represented in the reference, should receive inconsistent votes and thus lower scores. The prediction scores of each cell for each cell type category were projected onto UMAP clusters. The hESC-derived interneurons were also classified by using the developing macaque and mouse brain snRNAseq data^37^, which were downloaded from cells.ucsc.edu, which yielded consistent results (data not shown).

### Characterization of cells from other in vitro derivation methods

To compare our hESC-derived interneurons to GABAergic cells made by other in vitro derivation methods, the published scRNAseq datasets were downloaded and processed as described in the original reports. The datasets from each method were processed independently. Read counts of cells from Close et al.^87^ were downloaded from the Supplementary file of GSE93593, clustered and visualized with Seurat standard pipeline. Raw data from Allison et al.^88^ was downloaded from the Supplementary file of GSE180132. Two replicates of Ascl1/Dlx2 transcription factor reprogramming experiments^88^ were integrated and clustered with Seurat CCA methods, as guided by the original report. Read counts of cells from the 3i directed differentiation method^88^ were downloaded and clustered with Seurat standard pipeline. The raw FASTQ data of cortex + ganglionic eminence (CTX-GE) fusion organoids from Samarasinghe et al.^90^ were downloaded from GSE165577, demultiplexed by Cell Ranger v4.0.0 (10x Genomics), and only healthy control samples were included in the analysis. Samples from Days 56, 70, and 100 were integrated with Seurat CCA methods, as in the original report^39^. The cell types and pseudo-developmental ages were characterized independently by using the human GE reference, as described above.

### snRNAseq analysis of post-transplant cells

Following transplantations with hESC-derived interneurons, mouse brain cortices from 2-3 mice per sample were dissected at 1 month post-transplantation (MPT; N=3), 3-4 MPT (N=3), 6-7 MPT (N=4), 12 MPT (N=2) and 18 MPT (N=2), for a total of 14 samples across 5 timepoints using 5 independent hESC-derived interneuron lots (Table 1). Enrichment for human nuclei was done using FANS, followed by capture, barcoding, library generation and sequencing, as described above. Nuclei were demultiplexed for each sample with Cell Ranger v4.0.0 (10x Genomics).

### In-silico identification of human nuclei from sequenced post-transplant samples

To filter out contaminating mouse cells, we built a reference using both human (GRCh38) and mouse (mm10) genomes for the alignments of reads from each post-transplant sample. We calculated the percentage of human cDNAs for each barcoded cell by identifying the reads that aligned to human and mouse chromosomes. We observed two sharp peaks for most post-transplant samples, with one peak centered at 0% human cDNA, representing mouse cells, and the other peak centered at 100% human cDNA, representing human cells. As a control, only one peak was observed when pure human or pure mouse samples were aligned this way (data not shown). A cutoff of ≥80% human cDNAs was applied, thus removing any contaminating mouse cells and yielding high-quality human cells for downstream analyses.

### Identification of common cell types across post-transplant samples from different ages

To identify common cell types across all the samples, regardless of cell lot or age post-transplant, we applied a Seurat CCA integration workflow to identify cell clusters. Human nuclei were filtered for quality by having at least 1000 expressed genes, at least 1000 unique reads, no more than 1% reads of mitochondrial genes and no more than 1% reads of ribosomal genes. The data were normalized by the ‘LogNormalize’ method and the mean expression of genes between cells were scaled to 0. The top 3000 genes were identified by ‘FindVariableFeatures’ with ‘vst’ selection method, and the top 30 PCs were used for integration. UMAP was used to visualize cell clusters and gene expression.

### Characterization of post-transplant cell types by prediction analysis compared to human brain

To objectively interpret transplanted cell identity in terms of major cell classes, we compared our human nuclei snRNAseq data to the human adult whole brain snRNAseq dataset^92^, which sampled over 100 anatomically distinct brain regions derived from telencephalon, diencephalon, midbrain, hindbrain, and anterior spinal cord of several adults. In this dataset, 30 neuronal and non-neuronal superclusters corresponding to major cell classes were defined, providing a diverse transcriptional reference to which transplanted cells were compared. Both the processed data (Loom files) of gene expression and annotations of cells were downloaded. The dataset was downsampled to about 3E5 cells, due to the computational capability of our server. All reference superclusters were preserved for the analysis, except the “splatter” supercluster, since this population was likely under-sampled and/or under-sequenced^92^, leading to unclear interpretation of cell identities. PCA was applied, and the UMAP models of both cell types and brain regions were constructed with the whole adult brain dataset using Seurat. Human nuclei from hESC-derived cell transplantations were queried against adult brain cell types and regions, assigning prediction scores from 0 to 1, based on transcriptional similarity to the reference dataset.

### Characterization of interneuron subclasses and subtypes after transplantation by prediction analysis compared to human cortical interneuron datasets

To interpret the identity of transplanted cells in terms of interneuron subclasses, we selected multiple recently-published human brain datasets for reference, focusing on the studies that used the 10x Genomics platform, to avoid platform bias in single cell sequencing. We used human reference subclasses annotated by the BRAIN Initiative Cell Census Network (BICCN)^9^ and the human primary motor cortex (M1) snRNAseq dataset^93^ from the Allen Brain Institute. Subsequently, the middle temporal gyrus (MTG) snRNA-seq dataset was also published^10^, with larger cell numbers and subclass annotations based on analysis of multiple cortical areas, including M1. The gene expression matrices and cell annotations were downloaded from cell.ucsc.org, and the models of cell types were built using Seurat. Human nuclei from hESC-derived cell transplantations were queried against the annotated adult human inhibitory interneuron subclasses, as defined in the M1 and MTG reference datasets. To identify interneuron subtypes in the hESC-derived cell grafts, we determined the proportion of cells in each 18 MPT cluster that had high transcriptional similarity (prediction score) to one or more of the 72 transcriptionally defined MGE- and CGE-derived human interneuron subtypes defined in the cell census atlas of the mammalian primary motor cortex^9^. The identity of the 18 MPT subtypes was then inferred based on the documented overlap between the matching endogenous human and mouse consensus clusters, defined through multimodal analyses^9^.

### Identification of upregulated gene signatures over time after transplantation

To preserve the age difference among samples, we pooled all the nuclei from pre- and post-transplantation samples (without Seurat CCA integration) and performed independent clustering and UMAP visualization. In this analysis, most samples are grouped together based on age, regardless of the manufacturing lot. However, a few samples appeared to have batch effects due to technical reasons (sorting, capture and/or library preparation). To remove the non-biological batch effects, we applied a fast mutual nearest neighbor correction (fastMNN) method^136^ to correct the batch effects of some samples, while maintaining the differences between the different age groups. Upregulated genes in different stages of maturation were identified by the “FindAllMarkers” method from Seurat. Enriched pathway and GO terms in each stage were identified by the “compareCluster” method from clusterProfiler package^137^. To identify the genes that are upregulated after cell transplantation, the regression analysis from Monocle 3^138^ was used. Enriched pathway and GO terms in each stage were identified by the “compareCluster” method from clusterProfiler package^137^. To identify the genes that are upregulated after cell transplantation, the regression analysis from Monocle 3^138^ was used. (without Seurat CCA integration) and performed independent clustering and UMAP visualization. In this analysis, most samples grouped together based on age, regardless of the manufacturing lot. However, a few samples appeared to have batch effects due to technical reasons (sorting, capture and/or library preparation). To remove the non-biological batch effects, we applied a fast mutual nearest neighbors correction (fastMNN) method^136^ to correct the batch effects of some samples, while maintaining the differences between the different age groups. Upregulated genes in different stages of maturation were identified by the “FindAllMarkers” method from Seurat. Enriched pathway and GO terms in each stage were identified by the “compareCluster” method from clusterProfiler package^137^. To identify the genes that are upregulated after cell transplantation, the regression analysis from Monocle 3^138^ was used.Enriched pathway and GO terms in each stage were identified by the “compareCluster” method from clusterProfiler package^137^. To identify the genes that are upregulated after cell transplantation, the regression analysis from Monocle 3^138^ was used.

### Pseudo-age prediction analysis compared to human brain development

Recent studies have captured the molecular progression of cortical lineages across the human life span^36,94^. The snRNAseq dataset from Velmeshev et al.^36^ integrated several recent publications, including human prefrontal cortex (PFC) development from Herring et al.^50^, comprising postmortem PFC samples from 26 individuals spanning fetal, neonatal, infancy, childhood, adolescence, and adult stages of development, providing a comprehensive reference of human cortical development. The processed human cortical development snRNAseq data were downloaded from cells.ucsc.org. (doi: https://doi.org/10.1101/2022.10.24.513555) Only the inhibitory interneurons were used as a reference, and the models of cell types and cortical development ages were built using Seurat, as described above. Cell identities and development ages were characterized using prediction score compared to this reference, similar to other analyses. To investigate specific pathways, the ‘AddModuleScore’ from Seurat was used to calculate the module expression score for each group of genes or gene families. The ‘RidgePlot’ from Seurat or the ggridges package was used to visualize the module expression scores.

### Interneuron subclass assignments prior to transplantation and after 30 DIV culture on primary mouse glial feeder cells

Classification of pre-transplant and 30 DIV samples was guided by clustering with 14 DPT and 1 MPT snRNAseq data from the same batch. The SCT normalization method from Seurat was used, Seurat CCA integration was applied, enabling efficient clustering between pre-transplant and corresponding post-transplant samples. Cell fates of the in vitro samples were thus assigned based on co-clustering with the 1 MPT cells. Prediction analyses based on human cortical reference data sets were done, as described above.

---

**Supplemental Figure 1.**
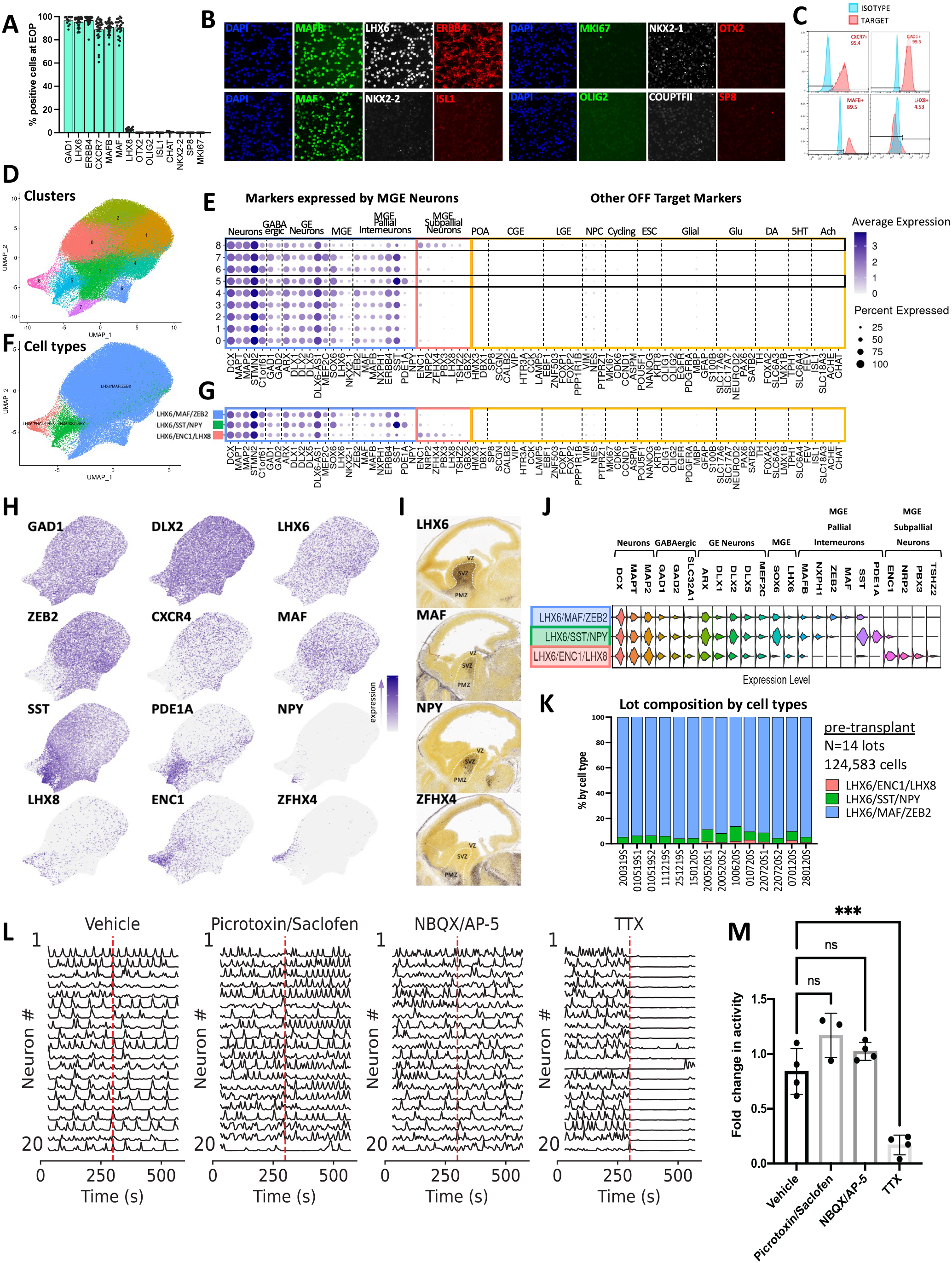
(Related to figure 1). Homogeneous expression of GABAergic MGE-type pallial interneuron markers and calcium transients prior to transplantation (A-C) Marker expression across ERBB4-sorted lots (N=14) was assessed using a combination of immunocytochemistry (LHX6, MAFB, MAF, NKX2-1, OTX2, OLIG2, ISL1, NKX2-2, SPI8, COUPTFII, MKI67), surface flow (ERBB4, CXCR7), intracellular flow (MAFB), and prime flow (*GAD1, LHX8, CHAT*) cytometry assays at the end of the differentiation process, post-thaw. (A) Quantification of key markers. Dots represent independent lots (each dot is an average of technical replicates), bar graphs shown mean ± SEM. (B, C) Representative examples of ICC (B) and flow (C) results for ON- and OFF- target markers. (D-K) Single cell RNA sequencing (scRNAseq) of 14 lots (total = 124,583 cells; median = 10,115 cells/lot). (D, F) Uniform Manifold Approximation and Projection (UMAP) visualization of cell clusters (D) and cell types (F). (E-H) Expression of key markers by clusters (E), cell types (G), and overlaid onto UMAP (H). Abbreviations: glutamatergic (Glu), dopaminergic (DA), serotonergic (5HT), cholinergic (Ach). (I) Expression of reference markers in E13.5 mouse brain by in situ hybridization (Allen Developing Mouse Brain Atlas, 2008). Ventricular zone (VZ), subventricular zone (SVZ) and pallidal mantle zone (PMZ). (J) Expression of shared and divergent markers across the three cell types. (K) Composition by cell type across 14 lots. (L, M) Effects of synaptic receptor antagonists on spontaneous calcium activity of hMGE-pINs after 17 DIV culture on primary mouse astrocytes. (L) Representative calcium traces show activity of 30 neurons before and 20 minutes after (dashed red line) application for each set of compounds. (M) Fold change in firing rate after application of vehicle (media), picrotoxin and saclofen (GABAA and GABAB receptor antagonists, respectively), NBQX and AP-5 (AMPA- and NMDA-receptor antagonists, respectively), and TTX (Na+ channel blocker). One way ANOVA with Bonferroni post-hoc analysis (ns-no significance, *p<0.05, **p<0.01, ***p<0.001).

**Supplemental Figure 2.**
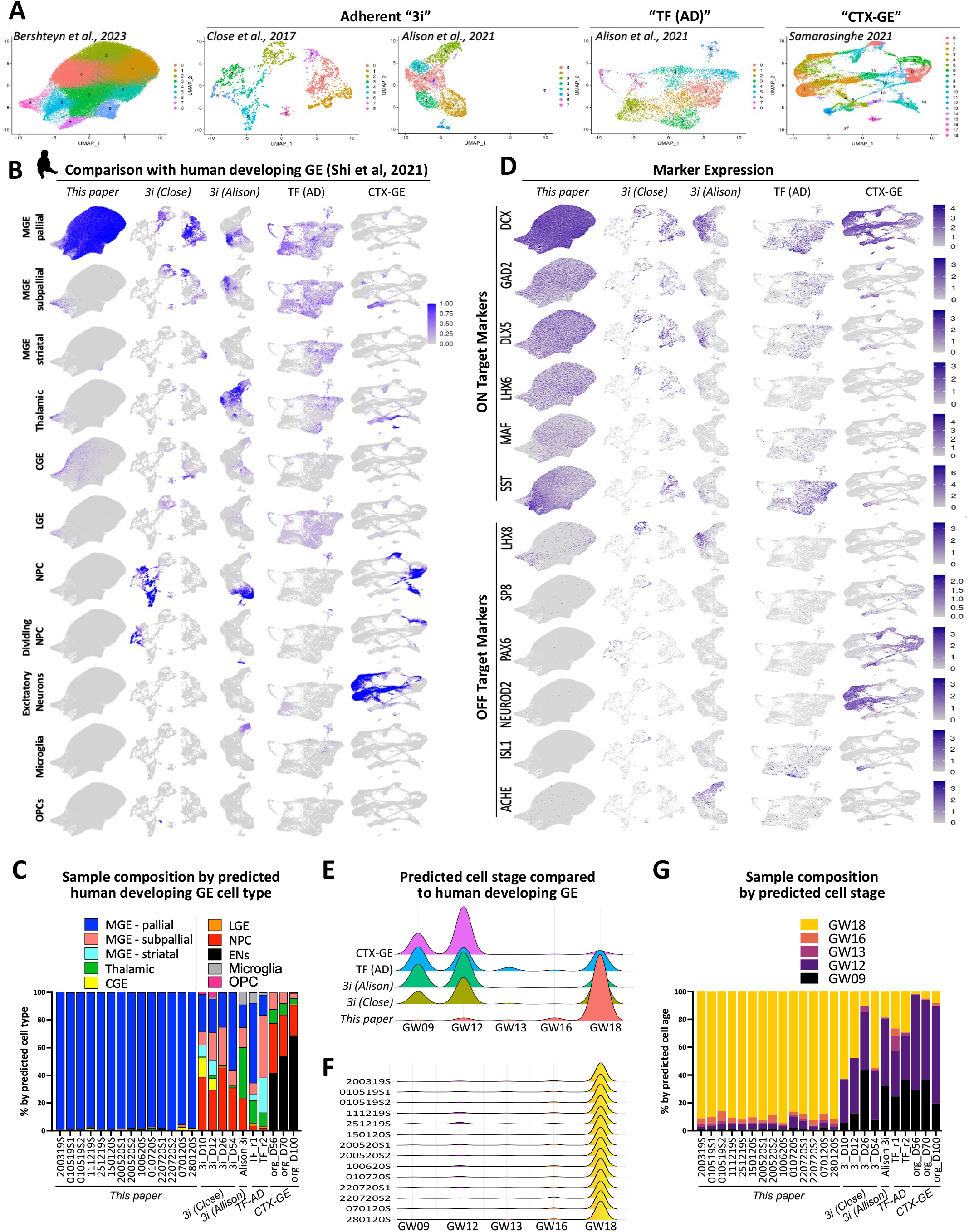
(Related to figure 1) High ON-target purity and stage-synchronicity of hMGE-pINs prior to transplantation. (A) Clustering analysis of published *in vitro* differentiation methods being compared. (B) Expression of key ON- and OFF-target markers across protocols. (C) Prediction analyses compared to human developing GE (gestational weeks, GW9-18)^35^ across protocols. (D) Sample composition based on predicted GE populations. (E-G) Transcriptional similarity to different stages of human GE development. (E) Predicted cell stage by protocol. (F) Predicted cell stage across hMGE-pIN lots. (G) Sample composition based on predicted cell stage.

**Supplemental Figure 3.**
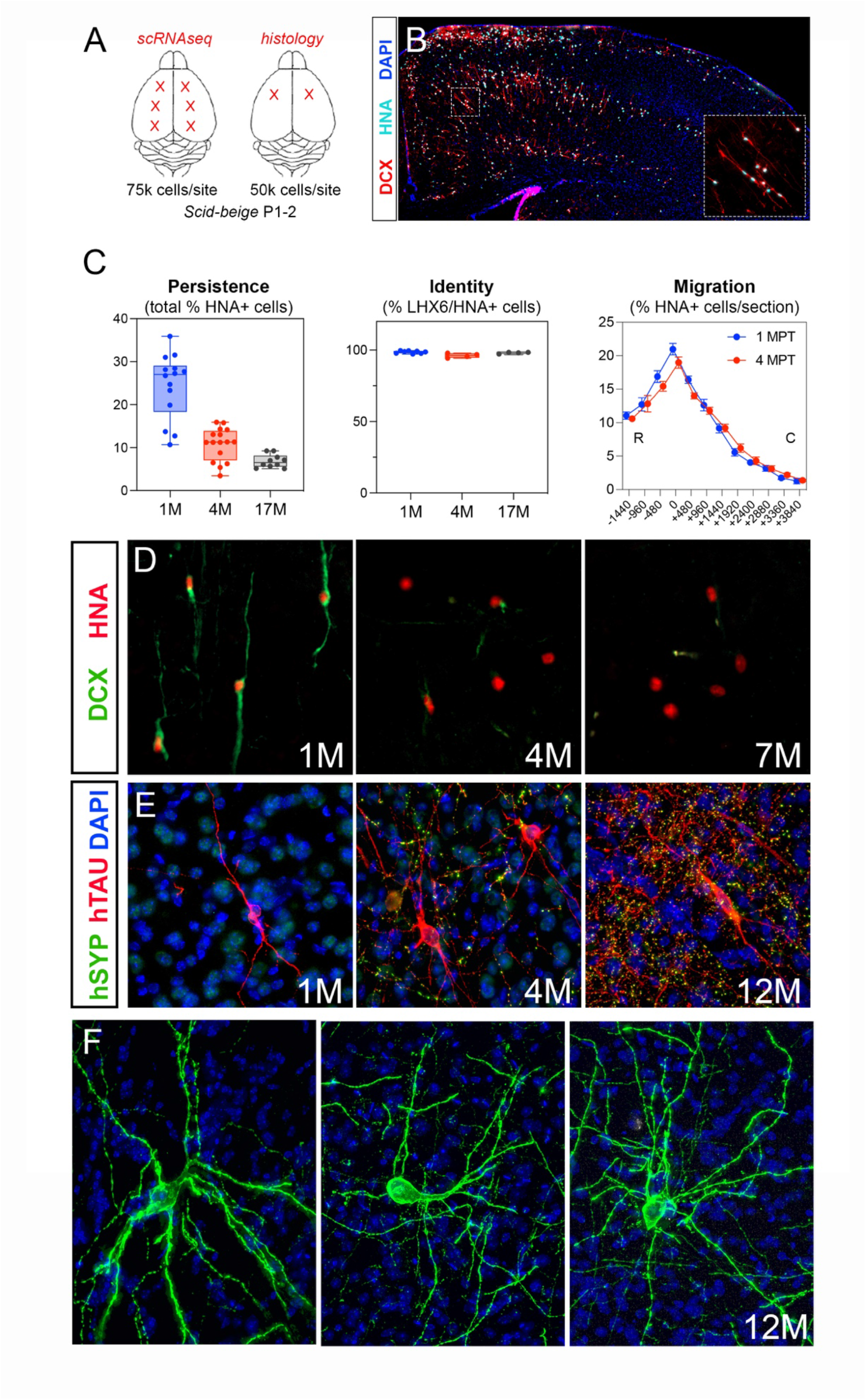
(Related to figure 1). Xenotransplantation of hMGE-pINs into mouse neonatal cortex. (A) Diagram showing transplantation sites and dosing in the neonatal cortex for scRNAseq and histological characterization. (B) Distribution of hMGE-pINs at 1 MPT, labeled with human-specific nuclear antigen (HNA), DCX, and LHX6 (coronal view). (C) Percentage of HNA+ cells persisting at 1, 4, and 17 MPT out of the number of cells that were injected (left), percentage of LHX6+HNA+ cells at 1, 4, and 17 MPT (middle), and relative rostral (R) to caudal (C) distribution of HNA+ cells at 1 and 4 MPT (right) (N=13 and 14 hemispheres for 1 and 4 MPT, respectively). (D) Expression of DCX at 1, 4, and 7 MPT. (E) Expression of the human-specific synaptic marker, SYP (synaptophysin) at 1, 4, and 12 MPT. Human-specific TAU denotes neuronal bodies, dendrites, and axons. (F) Examples of hMGE-pIN morphologies at 12 MPT. Membrane-bound GFP expression on the transplanted cells highlight their diverse morphological features as they mature in the host environment.

**Supplemental Figure 4.**
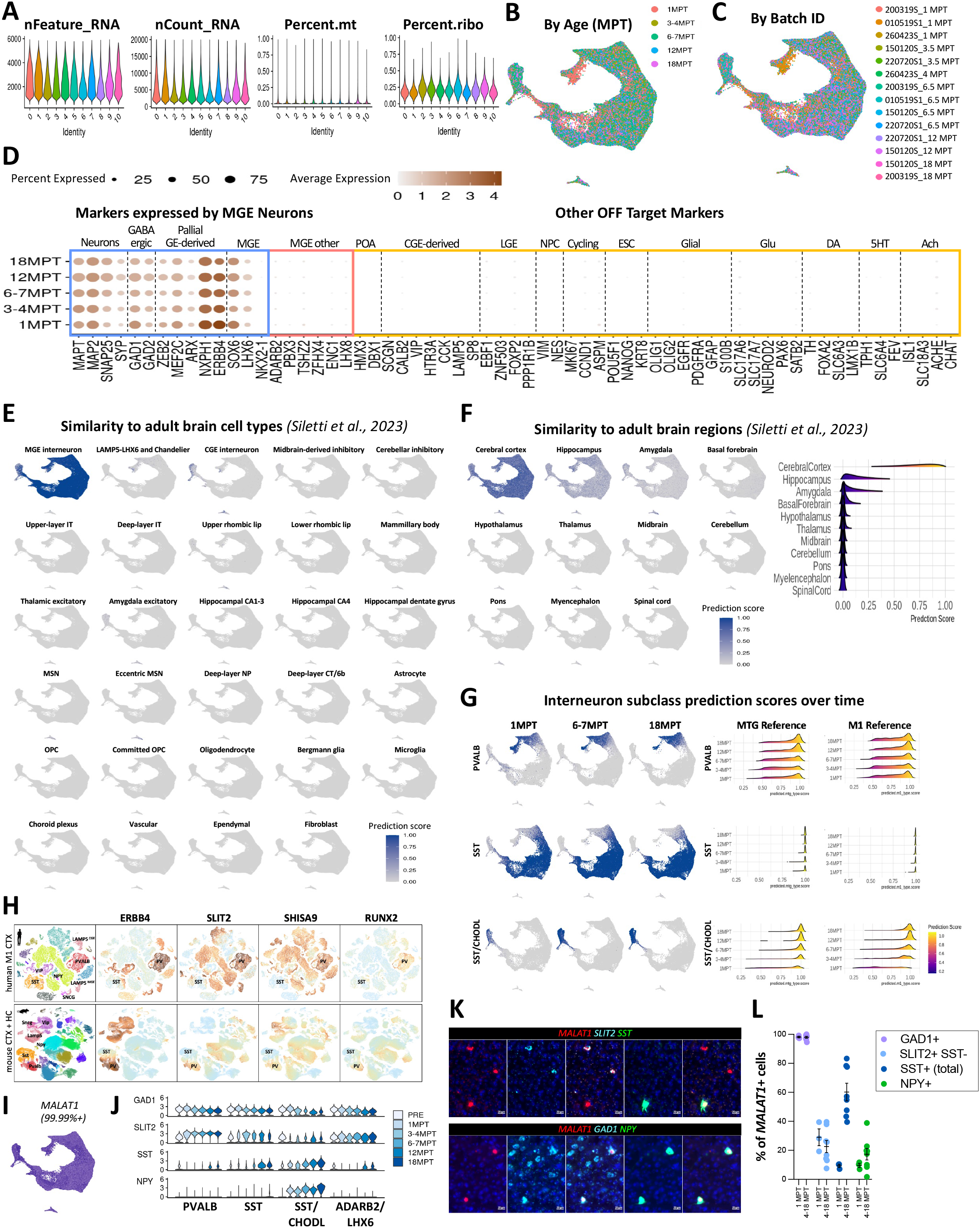
(Related to figure 1) Additional characterization of post-transplant cell identity. (A) Quality metrics of post-transplant cell clusters, including number of unique genes detected (nFeature), number of total RNA molecules (nCount), percentage of mitochondrial and ribosomal genes. (B, C) UMAP colored by Age (B) and by batch ID (C). (D) Expression of key genes over time. (E, F) Prediction scores projected onto the integrated UMAPs based on comparisons to the listed human adult brain^92^ cell types (E) and brain regions (F). The ridge plot on the right panel in F shows distribution of prediction scores for the transplanted population compared to each reference brain region. (G) Prediction scores projected onto the integrated UMAPs based on comparisons to endogenous human PVALB, SST and SST/CHODL subclasses, separated by age. The ridge plots show distribution of prediction scores for each subclass over time compared to adult human MTG^10^ (middle panels) and M1^93^ (right panels) cortical reference datasets. (H) Expression of several conserved markers (*ERBB4, SLIT2, SHISA9, RUNX2*) enriched in endogenous PVALB compared to SST INs in the adult human M1 cortex and mouse cortex + hippocampus (adapted from the Allen Brain Map: Cell Types Database). (I) *MALAT1* mRNA is expressed in 99.99% of post-transplant human cells, regardless of subclass or age. (J) Expression of *GAD1, SLIT2, SST*, and *NPY* transcripts by subclass over time from snRNAseq analysis. (K) Representative fluorescence in situ hybridization (FISH) images showing expression of listed targets in transplanted human cells in the mouse cortex. Note that the *GAD1* probe, although designed against human mRNA sequence, also recognizes endogenous mouse *Gad1*. (L) Quantification of human *MALAT1+* cells co-expressing listed targets. Each dot is from a different hemisphere, after averaging technical replicates (4 sections each). 1 MPT (n=3 hemi, 2 cell lots), 4-18 MPT (n=8 hemi, 3 cell lots). Lines show mean ± SEM.

**Supplemental Figure 5.**
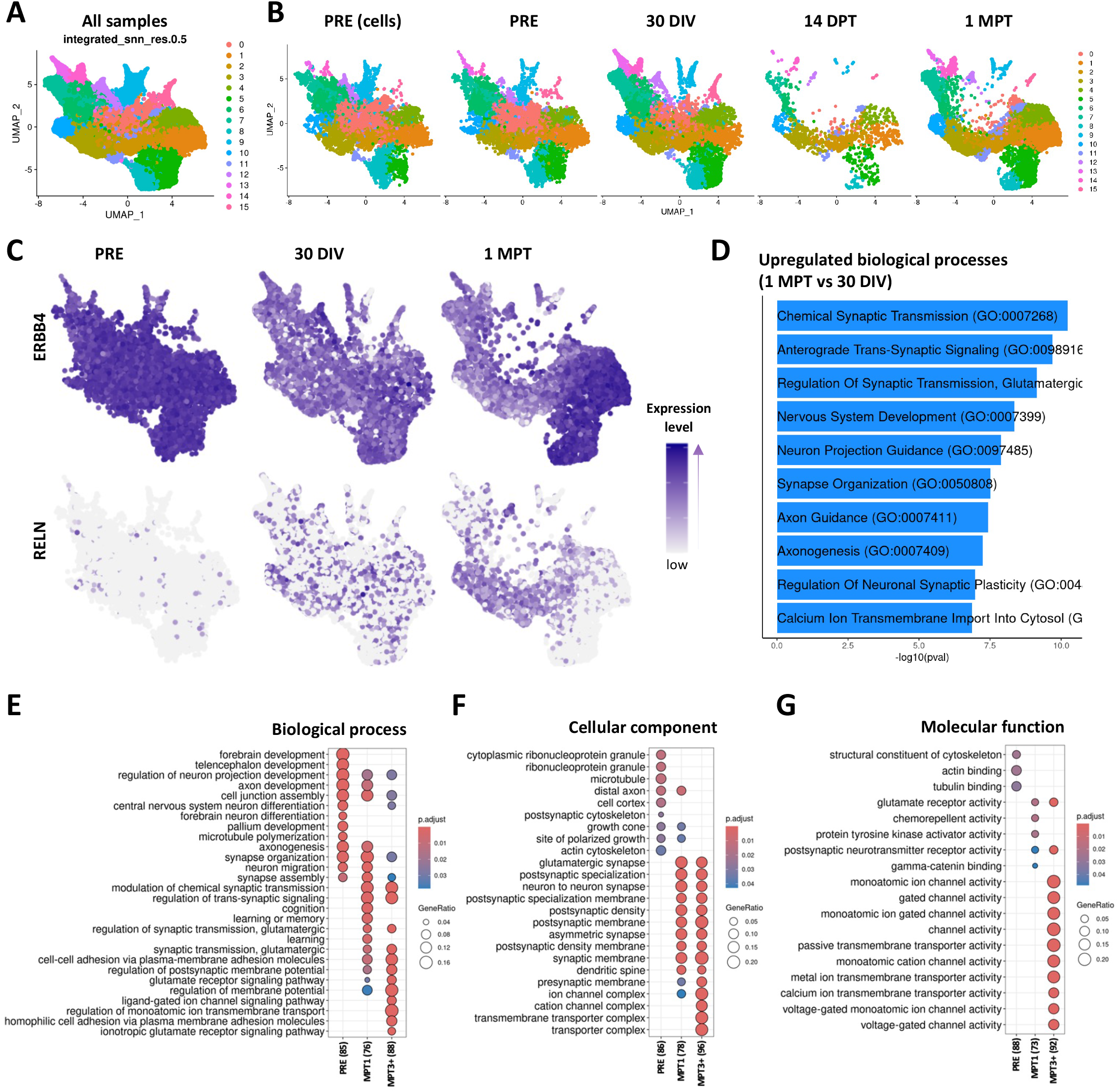
(Related to figures 2 and 3) Gene expression patterns under different conditions and at different stages post-transplant. (A, B) Integrated clustering analysis of lot 010519S1 sequenced prior to transplantation (PRE cells and PRE nuclei), after 30 days *in vitro* (DIV; nuclei), 14 DPT (nuclei) and 1 MPT (nuclei), colored by cluster (A) and separated by sample (B). (C) Expression of *ERBB4* (enriched in PVALB subclass) and *RELN* (enriched in SST subclass) in PRE, 30 DIV and 1 MPT nuclei samples. (D) GO term analysis for biological processes enriched in subclass-specific genes that are upregulated in 1 MPT vs 30 DIV samples. (E-G) GO term analysis for biological processes (E), cellular components (F), and molecular functions (G) enriched at each of the main transcriptional states: PRE, 1 MPT, and 3+ MPT.

**Supplementary Figure 6.**
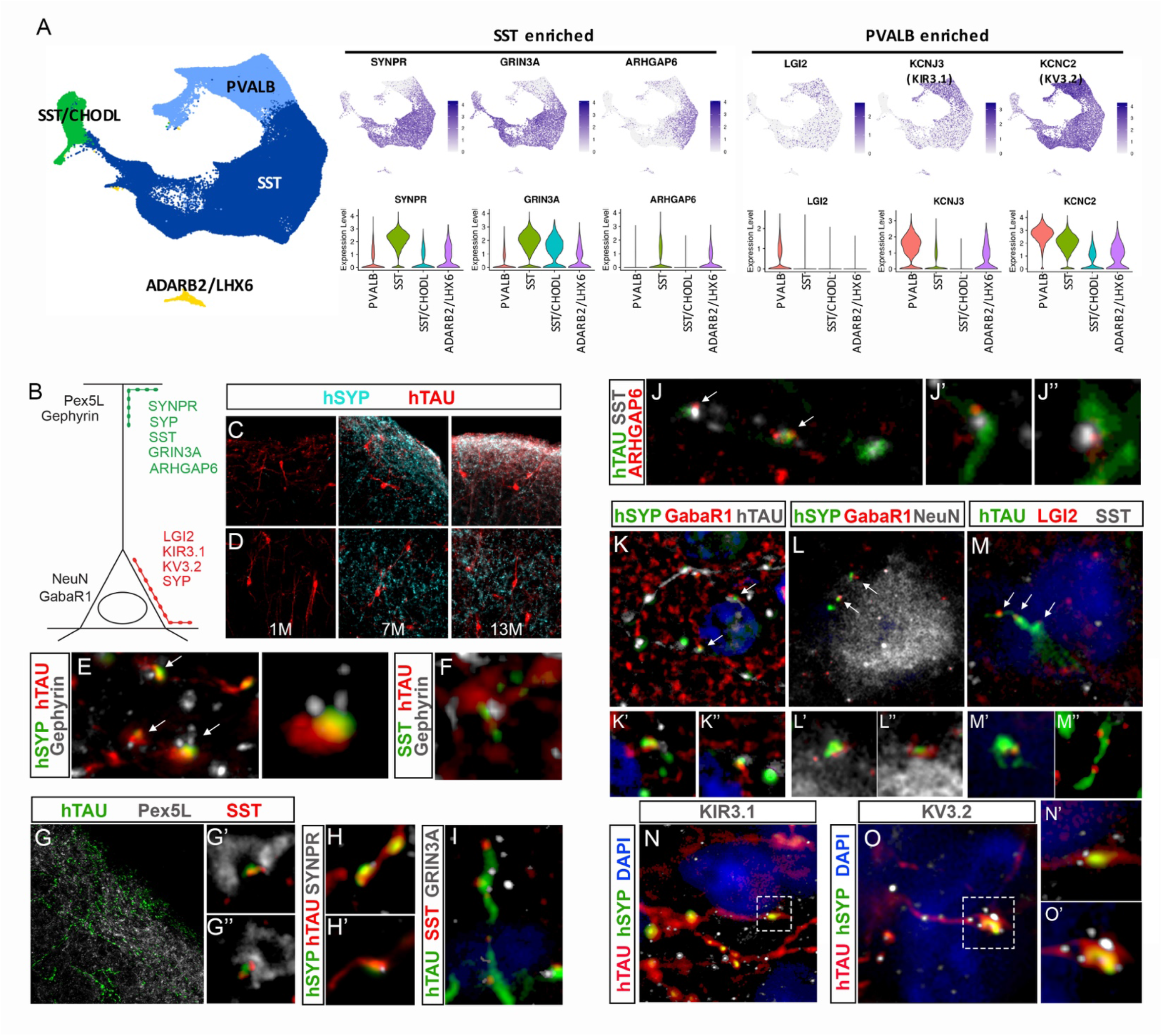
(Related to figure 6) hMGE-pINs exhibit subclass-specific synaptic connectivity features in the host mouse cortex. (A) Subtype-specific expression of synaptic and ion channel markers by snRNAseq. (B) Diagram showing a pyramidal projection neuron with dendritic (Pex5L) and somatic (NeuN), as well as presynaptic (Gephyrin and GabaR1) marker expression. Postsynaptic and axonal markers associated with SST (green) and PVALB (red) subclasses are illustrated collocalizing with human- specific TAU and SYP. (C-D) Increased cortical expression of the human-specific synaptophysin (SYP) in layers I-II (C) and IV (D) at 1, 7, and 13 MPT. (E, F) Examples of inhibitory synapses found in layer I composed of presynaptic human SYP and SST and postsynaptic Gephyrin (arrows). (G) SST-expressing human axonal terminals are intimately intertwined with Pex5L+ distal dendrites of pyramidal cells (G’, G” show higher magnifications with SST expression). (H, H’) Representative images showing co-expression of human-specific SYP and synaptoporin (SYPR) in axonal terminals in layers I/II. (I, J) Examples of human axonal processes in layer I co-expressing SST, the glutamate ionotropic receptor, GRIN3A (H), and the SST1-cluster marker, ARHGAP6 (arrows in J; additional examples are shown in J’ and J”). (K-M) Human axonal terminals show perisomatic synaptic patterns associated with PVALB cells in layer IV. Examples of perisomatic synapses formed between presynaptic human SYP (K, L) and LGI2 (M), and host cell postsynaptic GabaR1 (arrows). Bottom panels (K’-M”) correspond to additional examples for each marker combination at higher magnification. NeuN labels neuronal cell bodies (L). (N, O) Representative images showing perisomatic human axonal terminals co-expressing the PVALB-enriched potassium channels, KIR3.1 (encoded by *KCNJ3*) and KV3.2 (encoded by *KCNC2*) (N and O, respectively). Insets (N’ and O’) correspond to higher magnification of their respective images. Samples were obtained at > 7 MPT.

**Supplementary Figure 7.**
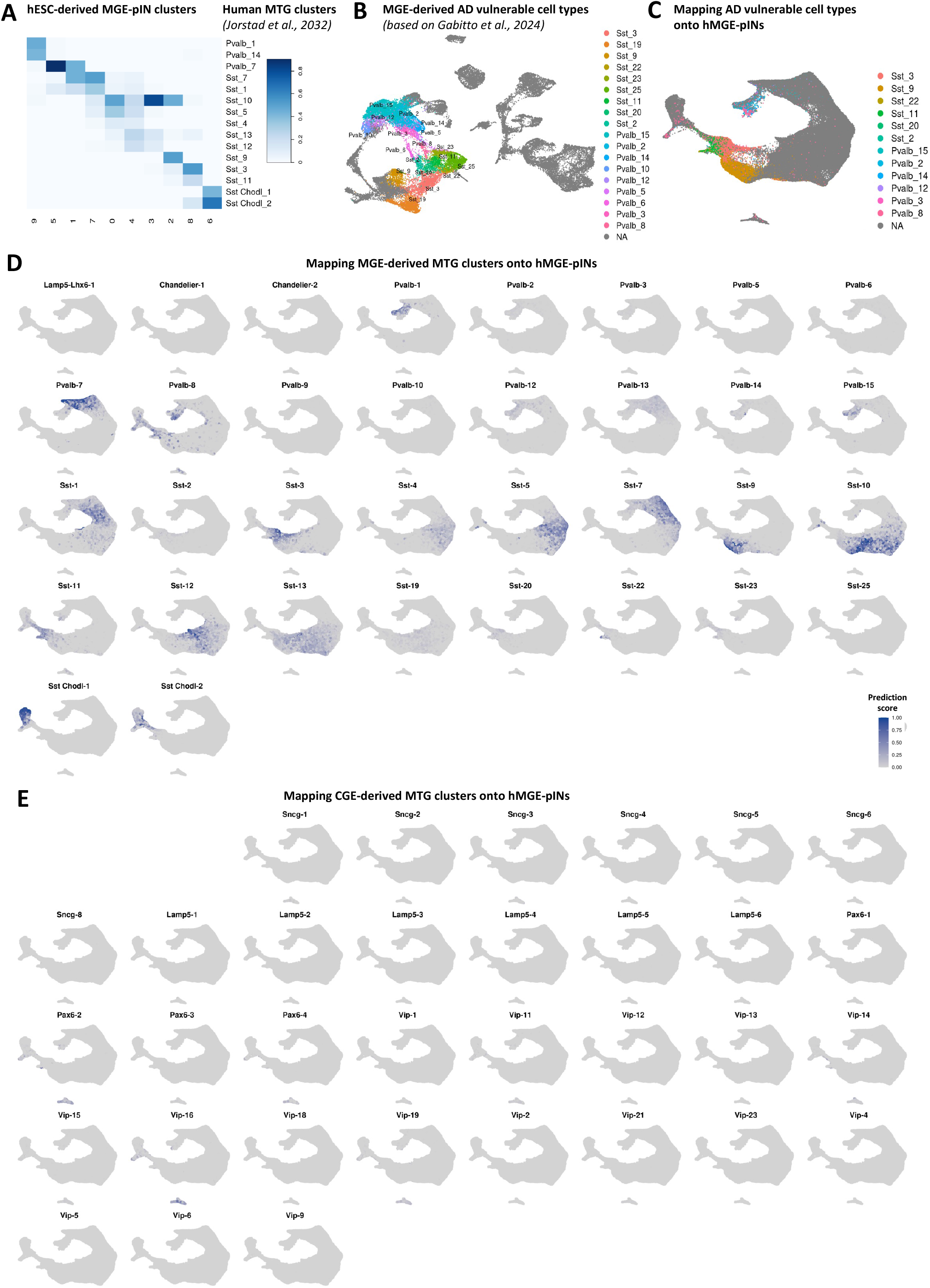
(Related to figure 6) AD vulnerable interneuron subtypes present in hMGE-pINs. (A) Distributions of predicted cell types for each 1-18 MPT cluster (columns) compared to endogenous human adult interneuron transcriptional clusters (rows) defined in the middle temporal gyrus (MTG)^10^. (B) The re-clustering of endogenous human MTG interneuron clusters, with colors representing the MGE-derived populations that were found to be significantly depleted in AD^10,106^. (C) Mapping AD vulnerable cell types from Gabitto et al.^106^ to hMGE-pINs. (D, E) Mapping all the MGE (D)- and CGE (E)-derived subtypes defined in MTG to hMGE-pINs based on transcriptional similarity (prediction scores).

