## Supplementary figures and images for "Human stem cell-derived GABAergic interneuron development reveals early emergence of subtype diversity followed by gradual electrochemical maturation"

### Supplemental Table 1

**Table 1. Summary of cell lots and data used for in vitro and in vivo sc/snRNAseq analyses**

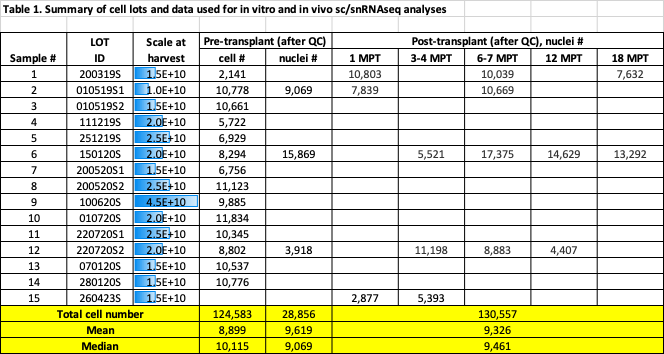
